# Racial and ethnic imbalance in neuroscience reference lists and intersections with gender

**DOI:** 10.1101/2020.10.12.336230

**Authors:** Maxwell A. Bertolero, Jordan D. Dworkin, Sophia U. David, Claudia López Lloreda, Pragya Srivastava, Jennifer Stiso, Dale Zhou, Kafui Dzirasa, Damien A. Fair, Antonia N. Kaczkurkin, Bianca Jones Marlin, Daphna Shohamy, Lucina Q. Uddin, Perry Zurn, Danielle S. Bassett

## Abstract

Discrimination against racial and ethnic minority groups exists in the academy, and the associated biases impact hiring and promotion, publication rates, grant funding, and awards. Precisely how racial and ethnic bias impacts the manner in which the scientific community engages with the ideas of academics in minority groups has yet to be fully elucidated. Citations are a marker of such community engagement, as well as a currency used to attain career milestones. Here we assess the extent and drivers of racial and ethnic imbalance in the reference lists of papers published in five top neuroscience journals over the last 25 years. We find that reference lists tend to include more papers with a White person as first and last author than would be expected if race and ethnicity were unrelated to referencing. We show that this imbalance is driven largely by the citation practices of White authors, and is increasing over time even as the field diversifies. To further explain our findings, we examine co-authorship networks and find that while the network has become markedly more integrated in general, the current degree of segregation by race/ethnicity is greater now than it has been in the past. Citing further from oneself on the network is associated with greater balance, but White authors’ preferential citation of White authors remains even at high levels of network exploration. We also quantify the effects of intersecting identities, determining the relative costs of gender and race/ethnicity, and their combination in women of color. Our findings represent a call to scientists and journal editors of all disciplines to consider the ethics of citation practices, and actions to be taken in support of an equitable future.

## Introduction

Race and ethnicity are socio-political categories that, while purporting to merely reflect physical or cultural differences, ultimately structure human groups and the social hierarchies among them. As techniques of controlling and governing those social hierarchies, racism and ethnocentrism are insidious and pervasive (1, 2). Beyond being ethically unjust, racism has a marked influence on mental and physical health (3). Racial and ethnic discrimination exists in academia as much as in any other sphere of our society and culture (4–12). Racial and ethnic bias impacts the promotion and retention of academics (5, 6), as well as grant funding and awards (7–10). Racial and ethnic bias also impacts the younger generation of scientists, limiting student access to prospective faculty mentors (11) and explaining the wide gap between the number of minority doctoral students graduated and the number of such students hired as faculty (12). Even when faculty of color are hired, racial and ethnic disparities exist in the number of scholarly publications, the number of citations they accrue, and the journal tier in which they appear (7).

Racial and ethnic disparities can arise from the explicit and implicit bias common in STEM fields (13, 14). Explicit bias refers to consciously held or expressed prejudice against a particular racial or ethnic group (15); implicit bias refers to subconsciously harbored discriminatory attitudes towards a particular racial or ethnic group that can result in prejudicial speech and social behaviors (16). Implicit bias can manifest in response to someone’s known race or ethnicity, or in response to someone’s appearance or name which is assumed (either correctly or incorrectly) to be an indicator of their race or ethnicity (17, 18). The same types of bias exist against minority genders in STEM, thus underscoring the need to understand intersecting identities (19) and their impact on academic success (20).

Addressing racial and ethnic disparities in academia is a challenge that requires concerted effort across many domains (21–25). It is true that marked responsibility lies in the hands of those in positions of power, such as journal editors, society presidents, institutional leaders, conference organizers, and those serving on committees to select award recipients, determine promotions, and grant tenure (26, 27). Yet there exists one tool that every scientist of any stature can utilize for good or ill: the reference list of their paper. Citations are a type of currency in academia (28), used to obtain career advancement in the form of jobs, tenure, promotion, grants, or other academic opportunities. They are also a record of science: the questions that have been and are being asked, and the answers of greatest import and value. Beyond impersonal records and goods of exchange, citations are often used and interpreted as inherently social acts, indicating the value that one human scientist places on (the work of) another human scientist. Citations can be distributed equally among all academics; or they can be distributed inequitably, actively excluding some people groups; alternatively, they can be distributed in a manner that actively favors people groups that have and continue to be adversely affected by histories, systems, and structures of inequality.

Recent work in several fields of science has identified a bias in citation practices such that papers from women, who are gender minorities in STEM, are under-cited relative to the number of such papers in the field (29–33). Tools to raise awareness and mitigate disparity are increasingly being developed and deployed (34–36). Here we seek to determine whether racial and ethnic imbalance exists in the reference lists of papers published in the field of neuroscience. Moreover, we go beyond a simple count and seek to explain any observed racial and ethnic imbalance in terms of the author’s own race or ethnicity, co-authorship networks, and the year in which papers were published. As we progress through these investigations, we pay special attention to how citation practices might or might not track the increasing diversification of the field, with implications for emerging challenges and potential solutions. Finally, we address the impact of intersecting identities, quantifying how gender and racial or ethnic imbalances combine and compound.

For this study, we examine articles published in five top neuroscience journals since 1995. Within this pool of articles, we obtain probabilistic estimates of authors’ presumed race based on their first and last names, find connections between citing and cited papers, locate and remove self-citations, and study the links between authors’ race/ethnicity and whether they are under-cited or whether they under-cite authors of a given race/ethnicity. Here and in what follows, we will use the phrase ‘author of color’ to refer to authors in the Asian, Black, and Hispanic racial/ethnic categories. It is important to note that the probabilistic models we use are a flawed approach for assessing something as personal, complex, and societally defined as race. However, both the self-identified and perceived race of a scholar can interact with a variety of individual and structural biases within the academy; to the extent that these models can accurately measure one or both of these characteristics, they can nontrivially capture the effects of such biases on authors of color. In this work, we do not seek, nor claim, to determine the ‘true’ race of any given author. Instead, we use aggregated probabilities of a racial/ethnic identity being associated with a given first/last name; we use these probabilities to reflect the cumulative impact of individual and structural biases based on name, appearance, and/or racial/ethnic identity.

Informed by recent evidence of gender imbalance in neuroscience reference lists (33), we test the following hypotheses: (1) The overall citation percentage of papers led by authors of color (defined here as those with people of color as first- and/or last-author) will be lower than expected given the characteristics of the paper that might be relevant to the number of times it is cited (e.g., the journal in which the paper was published, the seniority of the authors, and whether the paper was a review or an empirical article); (2) Under-citation of papers led by authors of color will occur to a greater extent within White-led reference lists; (3) Under-citation of papers led by authors of color will be decreasing over time, but at a slower rate within White-led reference lists; (4) Differences in under-citation between White-led and author-of-color-led reference lists will be partly explained by the structure of authors’ social networks; (5) Under-citation of papers led by authors of color will be greater for women of color than for men of color. In addressing hypotheses (1)-(4), we consider the two categories of White authors and authors of color; in addressing hypothesis (5), we consider the more granular categories of Asian, Black, Hispanic, and White authors. We acknowledge that we are using the term *author of color* broadly at both levels of granularity, and that future work would do well to study the distinct experiences of more specific people groups.

## Results

### Data collection and author race modeling

We extracted data from the Web of Science for research articles and reviews published in five top neuroscience journals since 1995. We selected the journals *Nature Neuroscience, Neuron, Brain, Journal of Neuroscience*, and *NeuroImage*, as they were reported by the Web of Science to have the highest Eigenfactor scores (37) among journals in the neuroscience category. In all, 63,677 articles were included in the dataset of citing/cited papers. See Methods for details on the procedures used to obtain and disambiguate full author names.

To assign author race or ethnicity, we used publicly available probabilistic databases and a deep neural network that learns the relationship between names and racial/ethnic categories (see Methods) (38, 39). We rely on given and surname data pulled from The Florida Voter Registration File from February 2017 and surname data pulled from the 2000 and 2010 Census (40). The 2000 and 2010 Census data uses the racial categories of American Indian/Native Alaskan, Asian/Pacific Islander, Black, Multi-Race, and White, and the ethnicity category of Hispanic. The Florida Voter Registration data uses the racial categories of Asian/Pacific Islander, non-Hispanic Black, and non-Hispanic White, and the ethnicity category of Hispanic. Here in the main text we provide results from the model built from the Florida Voter Registration data; this approach utilizes both first and last names, and has shown more balanced performance across racial/ethnic categories than other commonly used models. Using the model trained on these data, we can estimate the probability distribution across racial/ethnic categories based on each author’s first and last names. We study four categories to maximize statistical power for subsequent analyses (Asian, Black, Hispanic, White). Thus, for each author, we have four probabilities: one per racial/ethnic category. Sensitivity analyses using the United States Census data produced highly convergent results with those generated from the Florida Voter Registration data (see Extended Data).

### Trends in authorship

Across the articles in the sample, the proportion of articles with a person of color as first or last author increased between 1995 and 2019. Across these five journals, the overall probability of an article being published by either first- or last-authors of color increased from 47% in 1995 to 59% in 2019 (Figure 1a; Extended Data Figure 2a). To obtain a mean and confidence interval on the yearly increase, we performed 1000 bootstraps of the articles, summing over the probabilities associated with each racial/ethnic category. We found that the mean percent increase per year was 0.49% (95 CI:0.48%,0.51%). Within each journal, the average yearly increase in the probability of an article having a first or last author of color (C∪C) was 0.61% in *Brain*, 0.56% in *Journal of Neuroscience*, 0.47% in *Nature Neuroscience*, 0.59% in *NeuroImage*, and 0.57% in *Neuron* (Figure 1b,c; Extended Data Figure 2b,c; Extended Data Figure 3). We next use these data to determine whether citations are balanced or imbalanced relative to overall authorship proportions, before evaluating the structure and evolution of coauthorship networks.

**Figure 1.**
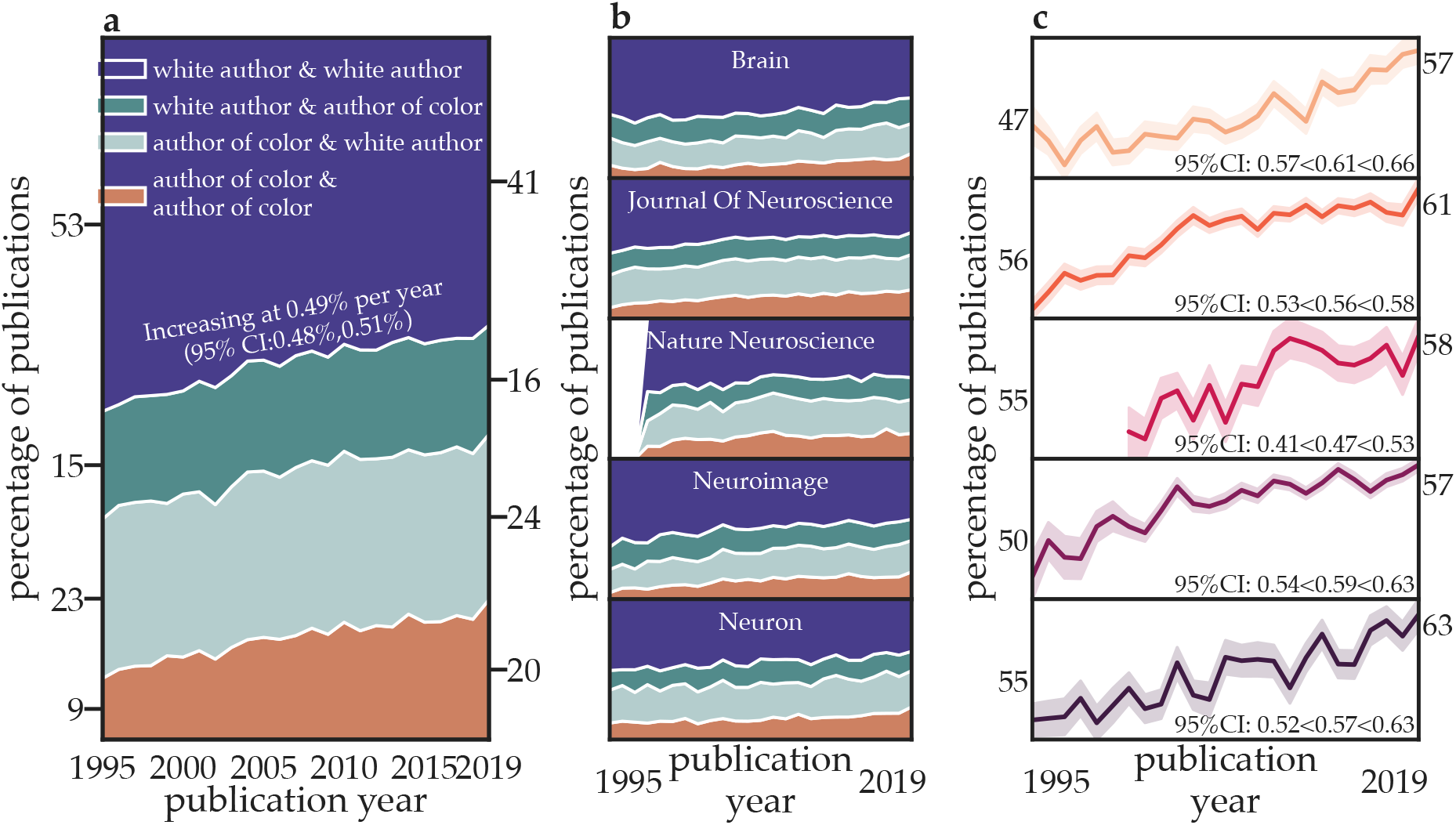
Racial and ethnic diversity is increasing within top neuroscience journals. **a** The percentage of papers published by authors of distinct racial/ethnic categories across the five journals studied from 1995 to 2019. **b** The percentage of papers published by authors of distinct racial/ethnic categories in each journal separately from 1995 to 2019. **c** The percentage of papers with a first and/or last author of color for each journal, for each year. Confidence intervals for the change in percent of papers with a first or last author of color (C∪C) for each year and for each journal were generated via bootstrapping (n=1000). Note: The journal *Nature Neuroscience* was not established until 1998. For complementary results using the Census model see Extended Data Figure 2, and for trends by racial/ethnic category, see Extended Data Figures 1 and 3.

### Citation imbalance relative to overall authorship proportions

To quantify citation behavior within neuroscience articles, we specifically examined the reference lists of papers published between 2009 and 2019 (*n* = 33,934). Thus, while all papers in the dataset were potential *cited* papers, references to *citing* papers refer only to those published since 2009. For each citing paper, we took the subset of its citations that had been published in one of the above five journals since 1995 and determined the probability that the cited first and last authors were from each of the four author categories. We removed self-citations (defined as cited papers for which either the first or last author of the citing paper was first-/last-author). We then calculated the probability that each of the cited papers fell into each of the four author categories: White author (first) & White author (last) (WW), White author & author of color (WC), author of color & White author (CW), and author of color & author of color (CC). Single-author papers by a White author were included in the WW category, and single-author papers by an author of color were included in the CC category.

For each citing paper, we compared the observed summed probabilities of citations within each category to the probabilities that would be expected if references were drawn uniformly at random from the pool of citable papers (Figure 2a, “Draw from literature”; Extended Data Figure 4). Each cited paper has four associated probabilities per author: one each for the racial/ethnic categories of Asian, Black, Hispanic, and White. We maintain all four probabilities, and simply sum over them across cited papers. To obtain the number of citations that would be expected under the assumption of random draws, we calculated the mean of author category probabilities among all papers published prior to the citing paper – thus representing the probabilities among the pool of papers that the authors could have cited – and multiplied them by the number of papers cited to match the observed summation of probabilities. Below, we expand on this naive measure to account for potential relationships between author categories and other relevant characteristics of cited papers, and to examine the relevance of co-authorship networks to citing behavior.

**Figure 2.**
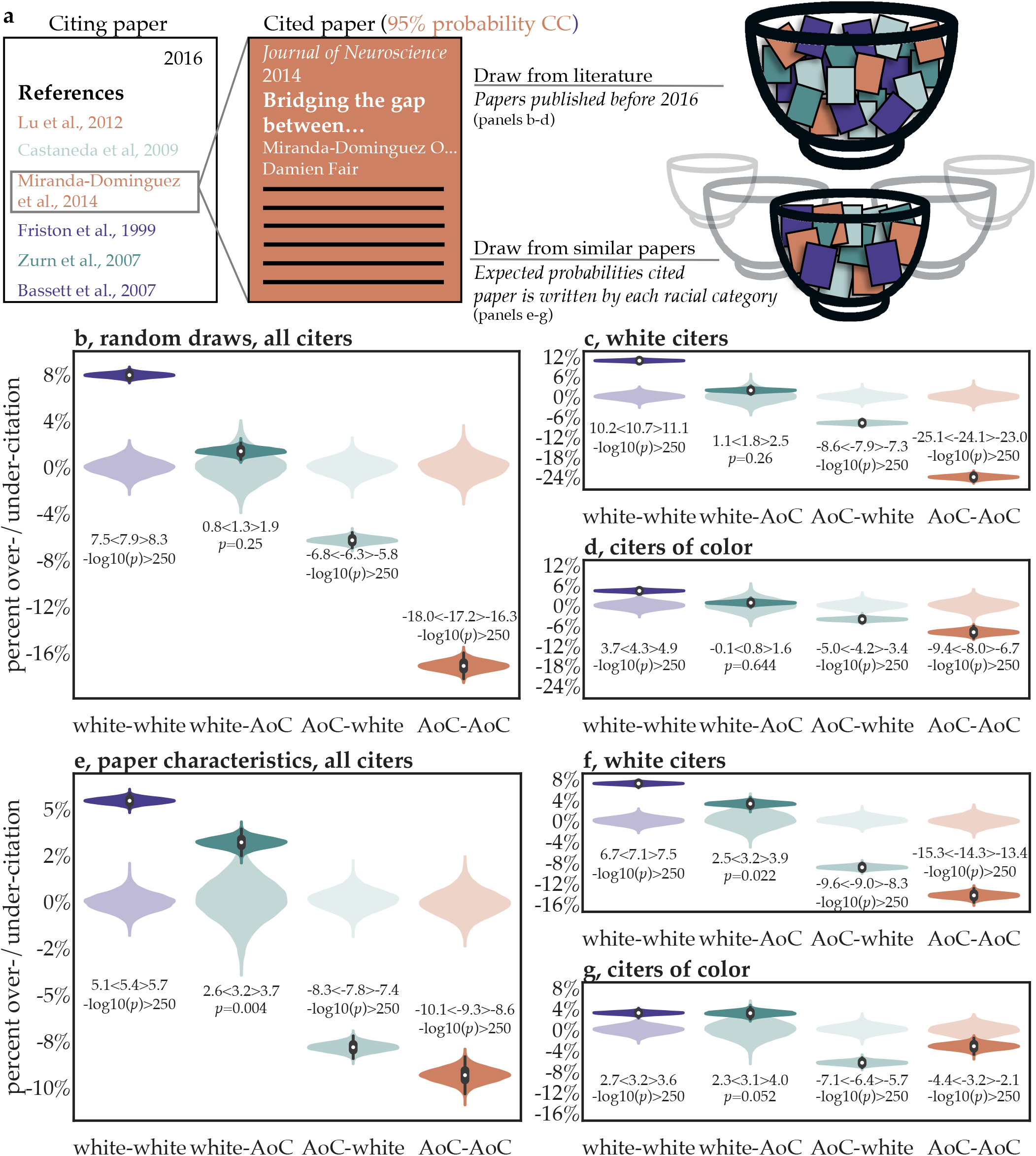
Over/under-citation of papers based on predicted author race and ethnicity. **a** In the random draws model, racial/ethnic category proportions in reference lists (permutations of White author and author of color (AoC)) are compared to the overall racial/ethnic category proportions of the existing literature. In the relevant characteristics model, author category proportions in reference lists are compared to author category proportions of similar articles by 1) year of publication, 2) journal of publication, 3) number of authors, 4) research article or review, 5) first and last author seniority, and 6) the location of the authors’ institution. **b** The over/under-citation of different author categories compared to their expected proportions under the random draws model are show in solid violin distributions; data for all citers are shown. Null distributions of over/under-citation are shown in transparent violins. **c,d** The same data as those in panel **b** but now separated by the racial/ethnic category of the citer: **c** White citers and **d** citers of color. **e** The over/under-citation of different author categories compared to their expected proportions under the relevant paper characteristics model; data for all citers are shown. **f,g** The same data as those in panel **e** but now separated by the racial/ethnic category of the citer: **f** White citers and **g** citers of color.

To estimate the amount of over/under-citation of each author category, we generated 10,000 bootstrap samples of citing papers. Each citing paper has an expected and observed citation percentage (the sum of probabilities) for each racial/ethnic category. Thus, in each bootstrap, the sums of observed and expected probabilities for each author racial/ethnic category are calculated across the sampled citing papers, each of which are normalized to sum to 1. The percentage over/under-citation relative to the expected proportion is the (observed % - expected %)/expected % (see Methods). For formal statistical inference, we construct a null distribution of over/under-citation percentages. To do so, we generate null “observed” citation counts for each paper by randomly re-drawing its citations from the pool of previously published papers. We then compare the resulting proportions to the expected proportions described above, yielding a zero-centered distribution of over/under-citation percentages that would be expected if the random draws assumption were true. Later sections account for additional characteristics of papers that affect citation; in those sections, a different null model is used for inference.

Of the 365,859 citations given between 2009 and 2019, WW papers received 50.4%, WC papers received 15.8%, CW papers received 22.0%, and CC papers received 11.7%. The expected proportions based on the racial/ethnic probabilities in the pool of citable papers were 46.7% for WW, 15.6% for WC, 23.5% for CW, and 14.1% for CC. By this measure, WW papers were cited 7.9% more than expected, WC papers were cited 1.3% more than expected, CW papers were cited 6.3 % less than expected, and CC papers were cited 17.2% less than expected (Figure 2b). These values correspond to WW papers receiving approximately 13530 more citations than expected, compared to roughly 770 more for WC papers, 5430 fewer for CW papers, and 8870 fewer for CC papers within references lists.

### The effect of authors’ race on citation behavior

By focusing the present analyses on the racial and ethnic composition of reference lists, we are able to investigate the racial/ethnic categories of the citing authors in addition to those of the cited authors. Here we compare the racial and ethnic composition of references within papers that had White first and last authors (WW) to those within papers that had a person of color as either first or last author (C∪C, comprising WC, CW, and CC papers).

After separating citing articles by author race, we find that the imbalance within reference lists shown previously is driven largely by the citation practices of WW teams (Figure 2c). Of the 206,729 citations given between 2009 and 2019 by WW papers, WW papers received 51.7%, compared to 15.9% for WC papers, 21.6% for CW papers, and 10.7% for CC papers. The expected proportions based on the pool of citable papers were 46.7% for WW, 15.6% for WC, 23.5% for CW, and 14.1% for CC. By this measure, WW papers were cited 10.7% more than expected, WC papers were cited 1.8% more than expected, CW papers were cited 7.9% less than expected, and CC papers were cited 24.1% less than expected. These values correspond to WW papers receiving approximately 10300 more citations than expected, compared to 580 more for WC papers, 3850 fewer for CW papers, and 7030 fewer for CC papers within WW reference lists.

Of the 159,130 citations given between 2009 and 2019 by C∪C teams, WW papers received 48.7%, compared to 15.8% for WC papers, 22.5% for CW papers, and 13.0% for CC papers. The expected proportions based on the pool of citable papers were 46.7% for WW, 15.6% for WC, 23.5% for CW, and 14.1% for CC. By this measure, WW papers were cited 4.3% more than expected, WC papers were cited 0.8% more than expected, CW papers were cited 4.2% less than expected, and CC papers were cited 8.0% less than expected (Figure 2d). These values correspond to WW papers receiving approximately 3190 more citations than expected, compared to 190 more for WC papers, 1570 less for CW papers, and 1810 less for CC papers within C∪C reference lists. The fact that C∪C teams do not overcite other C∪C teams suggests that citation behavior is not well-explained by outgroup bias.

### Citation imbalance after accounting for papers’ relevant characteristics

The comparison of citations to overall authorship proportions does not take into account other properties of published papers that may make them more or less likely to be cited by later scholarship. The potential relationships between author race/ethnicity and papers’ other characteristics make it difficult to isolate links between race/ethnicity and citation rates. To address this issue, we sought to model, and then account for, any relationships between race/ethnicity and paper characteristics. The features of a paper that we selected as being potentially relevant for citation were 1) year of publication, 2) journal of publication, 3) number of authors, 4) research article or review, 5) first and last author seniority, and 6) the location of the authors’ institution. For each paper, we have four values that estimate the probabilities that the first author is Asian, Black, Hispanic, or White, given their name, and four values that estimate these same probabilities for the last author. Thus, for example, if the probability that the first author is White is 0.50 and the probability that the last author is Black is 0.50, then the joint probability that the paper is authored by a White first author and a Black last author is 0.25. We use ridge regression – where the L2 regularization parameter has been determined by cross validation – to predict the probability of each of the 16 possible racial/ethnic category pairs as a function of the above characteristics. The process yields estimated author race/ethnicity pairing probabilities for each paper.

We then sought to compare the observed citation rates for each racial/ethnic category to those that would be expected if only paper characteristics were relevant to citation. Specifically, we estimate the expected racial/ethnic proportions across random draws from a narrow pool of papers highly similar to the cited paper (see Figure 2a, “Draw from similar papers”). For each citing paper, we generated null expected citation counts using the probabilities from the model that account for paper characteristics, and we generated null observed citation counts by pseudo-randomly citing based on those probabilities. This method generates an expected null distribution of over/under-citation rates that accounts for the possibility that papers from certain author categories are highly cited for reasons unrelated to their racial/ethnic category.

Summing up the number of cited papers from each category again gives us the observed citation rates, and summing up the estimated racial/ethnic category probabilities across all cited papers gives us new expected citation rates. Of the 365,859 citations given between 2009 and 2019, WW papers received 50.4%, compared to 15.8% for WC papers, 22.0% for CW papers, and 11.7% for CC papers. The expected probabilities after accounting for relevant paper characteristics of cited papers were 47.8 % for WW, 15.4% for WC, 23.9% for CW, and 12.9% for CC. By this measure, WW papers were cited 5.4% more than expected, WC papers were cited 3.2% more than expected, CW papers were cited 7.8% less than expected, and CC papers were cited 9.3% less than expected (Figure 2e; Extended Data Figure 4). These values correspond to WW papers receiving approximately 9490 more citations than expected, compared to 1790 more for WC papers, 6860 fewer for CW papers, and 4420 fewer for CC papers within all references lists. See Extended Data Figure 5 for similar results after accounting for different subfields of neuroscience.

After separating citing articles by author racial/ethnic category, we find that the imbalance within reference lists shown previously is driven largely by the citation practices of WW teams. Of the 206,729 citations given by WW teams between 2009 and 2019, WW papers received 51.7%, compared to 15.9% for WC papers, 21.6% for CW papers, and 10.7% for CC papers. The expected probabilities after accounting for relevant paper characteristics were 48.3% for WW, 15.4% for WC, 23.8% for CW, and 12.5% for CC. By this measure, WW papers were cited 7.1% more than expected, WC papers were cited 3.2% more than expected, CW papers were cited 9.0% less than expected, and CC papers were cited 14.3% less than expected (Figure 2f; Extended Data Figure 4f). These values correspond to WW papers receiving approximately 7090 more citations than expected, compared to 1030 more for WC papers, 4410 fewer for CW papers, and 3710 fewer for CC papers within WW references lists.

Of the 159,130 citations given by C∪C teams between 2009 and 2019, WW papers received 48.7%, compared to 15.8% for WC papers, 22.5% for CW papers, and 13.0% for CC papers. The expected probabilities after accounting for relevant paper characteristics were 47.2% for WW, 15.3% for WC, 24.1% for CW, and 13.4% for CC. By this measure, WW papers were cited 3.2% more than expected, WC papers were cited 3.1% more than expected, CW papers were cited 6.4% less than expected, and CC papers were cited 3.2% less than expected (Figure 2g; Extended Data Figure 4g). These values correspond to WW papers receiving approximately 2370 more citations than expected, compared to 760 more for WC papers, 2440 fewer for CW papers, and 690 fewer for CC papers within C∪C reference lists.

### Temporal trends in over/under-citation

We next sought to determine whether racial and ethnic imbalance in citations has changed as the field has become more diverse. Across 10,000 bootstraps, we calculate the slope of the linear relationship between the year of publication and the over/under-citation percentage for each racial/ethnic category. As above, we generate a null distribution for formal statistical inference by sampling the racial/ethnic probabilities from the model that incorporates salient paper characteristics. We found that the over-citation of WW papers is significantly increasing with time, both among White citers (Figure 6a) and among citers of color (Figure 3b); Extended Data Figure 6a,b). We also found that the under-citation of CC papers is significantly increasing in both White citers and citers of color (Figure 3a,b; Extended Data Figure 6a,b). Figure 6c-d compares the observed citation proportions over time with the proportions that would be expected under the paper characteristics model. The trends suggest that increasing under-citation of AoC-led teams is driven by stubbornly consistent (flat) citation proportions despite rapidly increasing expected citation proportions for AoC-led papers.

**Figure 3.**
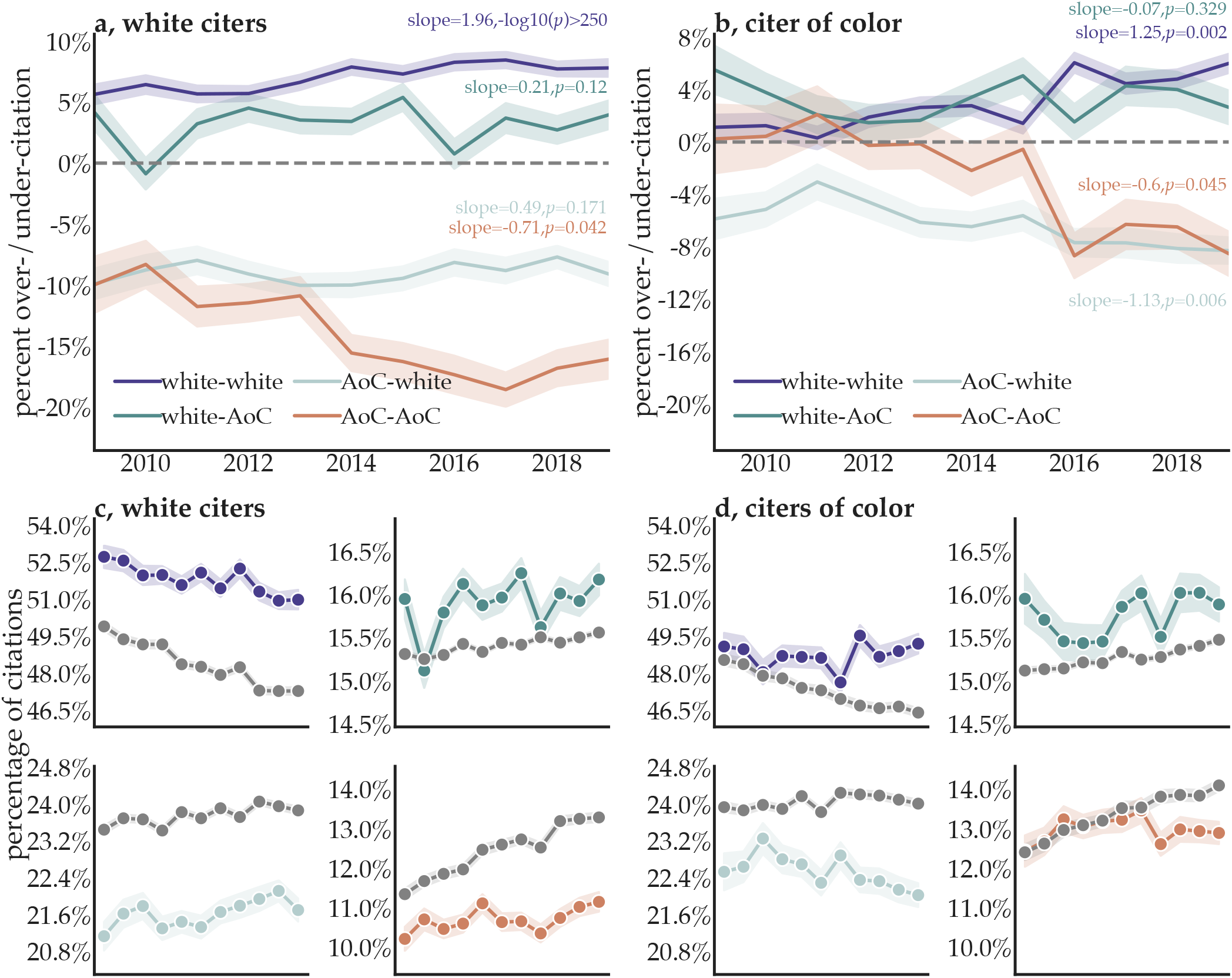
Temporal trends of over/under-citation. **a,b** The extent of over/under-citation across author racial/ethnic categories as a function of time, within WW (**a**) and C∪C (**b**) reference lists, under the paper characteristics model. The line represents over/under-citation within the literature in a given year. Shaded regions represent the 95% confidence interval of each over/under-citation estimate, calculated from 1,000 bootstrap resampling iterations. **c,d** Observed (colored) and expected (grey) citation proportions within WW (**c**) reference lists and C∪C (**d**) reference lists. Within each section, we show the observed and expected proportion of citations given by that group to WW papers (*top left*), WC papers (*top right*), CW papers (*bottom left*), and CC papers (*bottom right*). Points represent the estimated citation percentage as a function of year; shaded regions represent the 95% confidence interval of the observed citation rate, calculated from 1,000 bootstrap resampling iterations.

**Figure 4.**
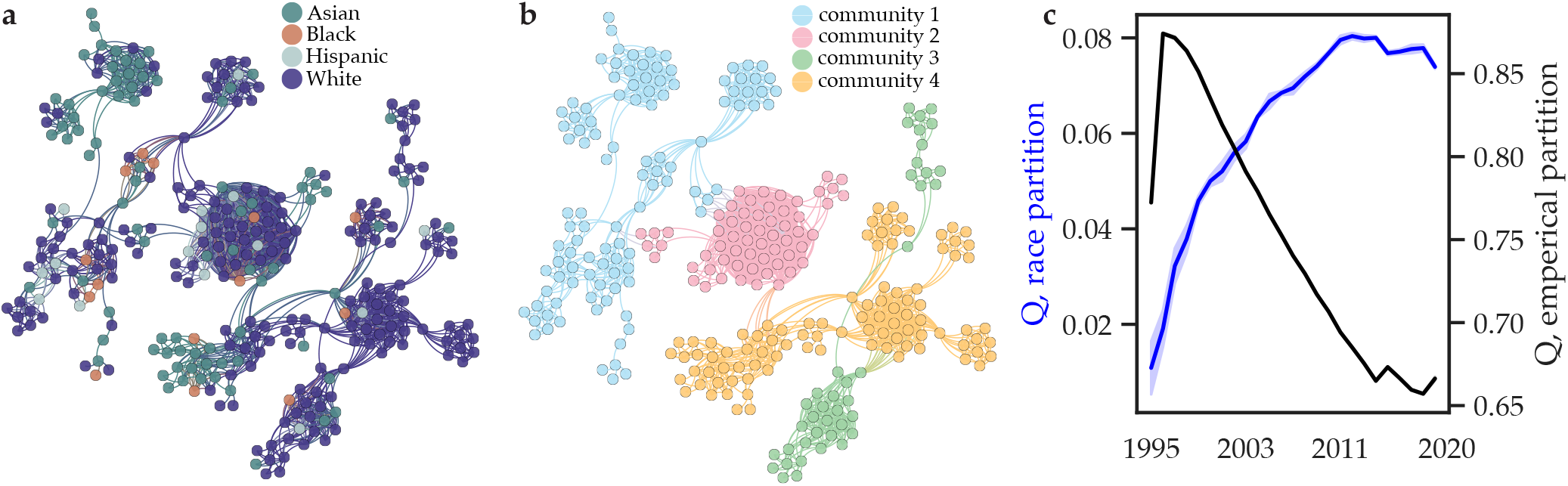
Racial and ethnic segregation in co-authorship networks is increasing despite greater field-wide collaboration. **a** The 2019 co-authorship network which we have colored according to the authors’ racial/ethnic category. **b** The same 2019 co-authorship network which we have colored according to a data-driven partition of authors into densely connected groups using the InfoMap community detection algorithm. **c** The modularity quality index, *Q*, evaluates the degree to which a partition of authors into groups explains the structure of the co-authorship network. While the data-driven partition consistently exhibits a higher Q than the racial/ethnic partition (**a,b**), here we sought to measure the temporal change in *Q*. For each year, we consider 100 partitions extracted by the Infomap algorithm (black) and 100 partitions that separate authors according to their racial/ethnic category (blue); for the latter, authors are assigned to a race/ethnicity based on the racial/ethnic category probabilities. Shaded areas indicate 95 percent confidence intervals.

**Figure 5.**
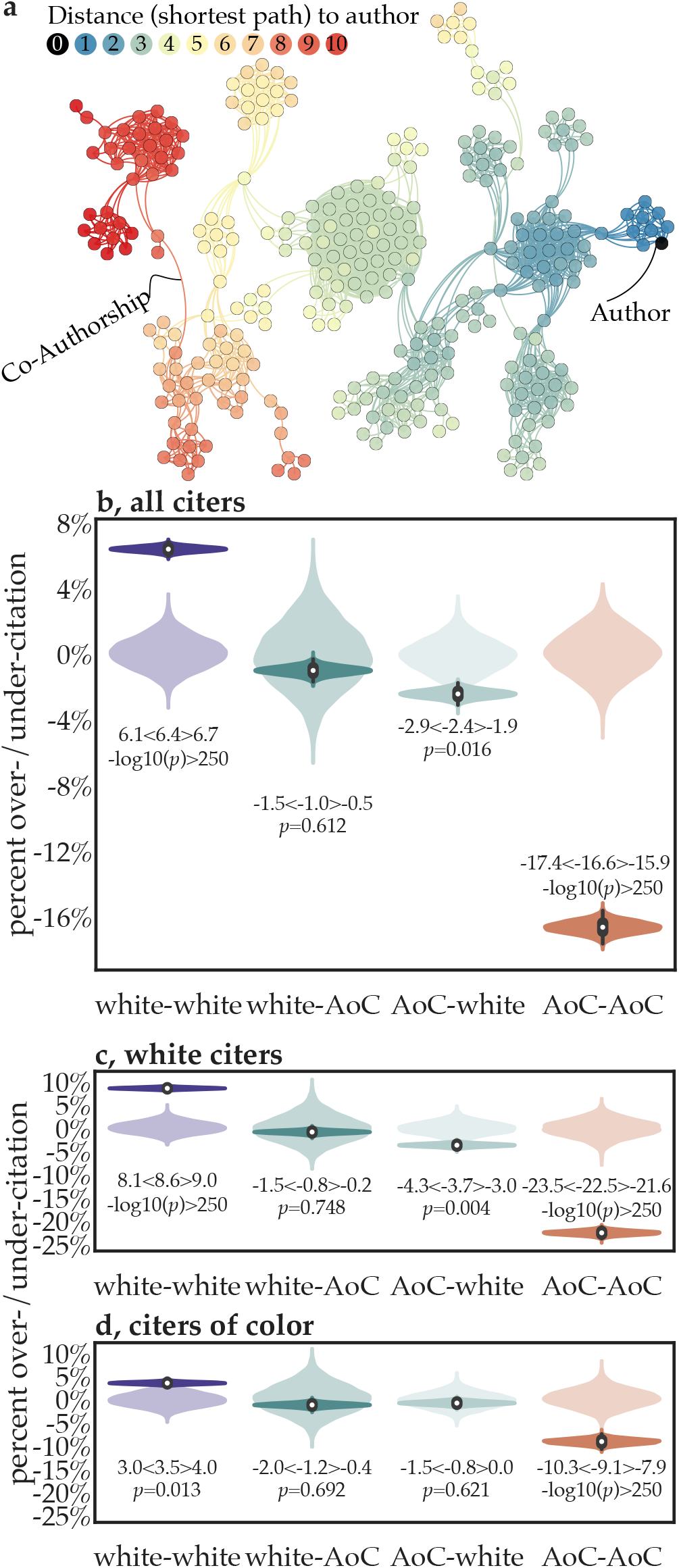
Over/under-citation of papers as a function of the distances between authors in the co-authorship network. **a** Shortest path distances in the co-authorship network from the citing author (black node) to other authors; node color indicates the length of the shortest path. **b** The over/under-citation of different author groups compared to their expected proportions under the shortest paths model for racial categories. In the lightly shaded violins, we show the null distributions under permutations of the author race/ethnicity. For each paper, we chose citing papers by randomly selecting authors that are the same distance away as the actual authors in the paper’s reference list. The same data as those in panel **b** but now separated out by the racial/ethnic category of the citer: **c** White citers and **d** citers of color.

**Figure 6.**
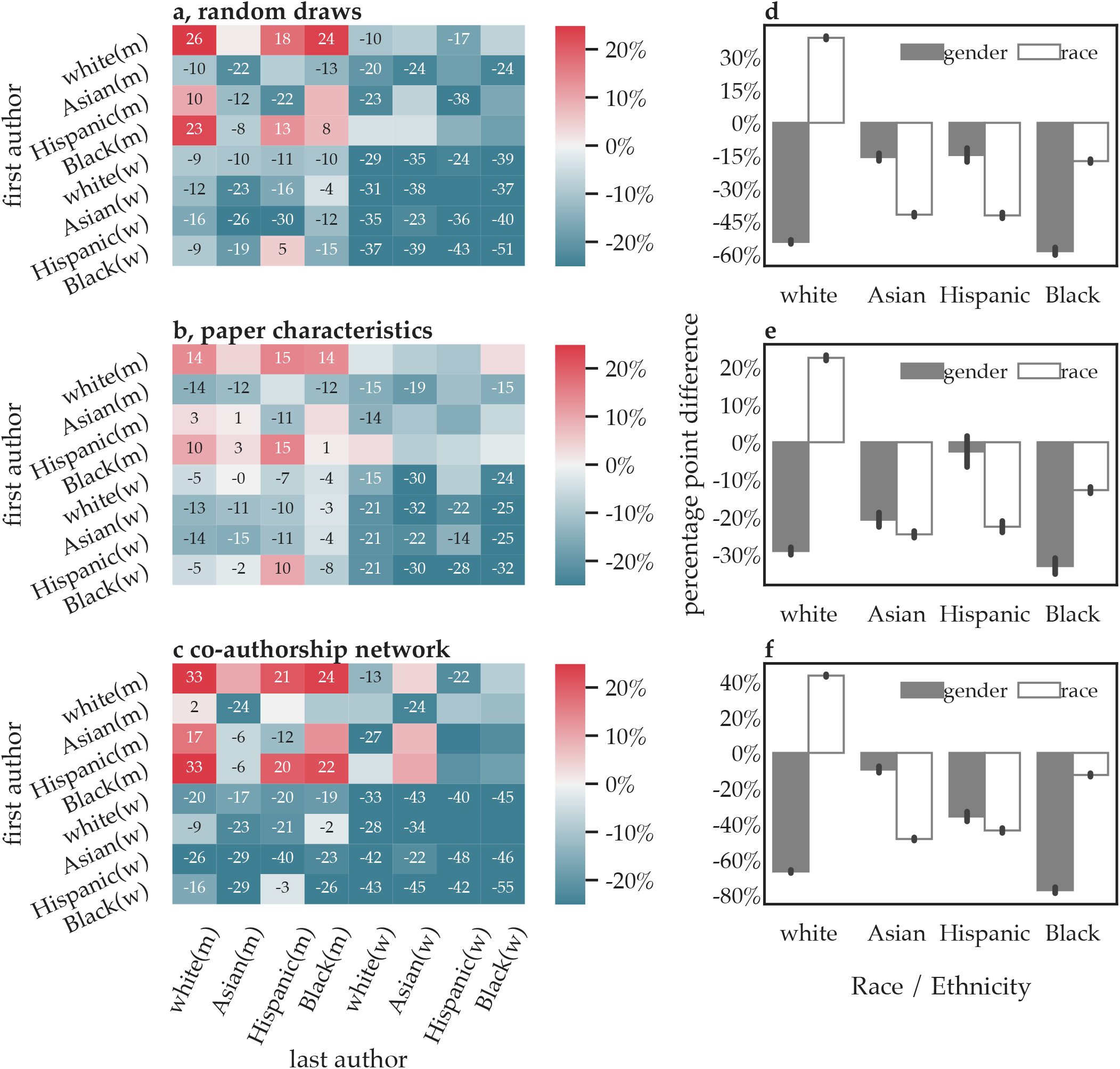
Citation costs at the intersection of gender and race/ethnicity. (**a-c**) We calculated the over/under-citation of papers for each combination of first author racial/ethnic and gender categories (y-axis) and last author racial/ethnic and gender categories (x-axis). Here, we compare the observed citation rates for each racial/ethnic and gender category to those that would be expected if base rates were defined by the random draws model (**a**), the paper characteristics model (**b**), or the shortest paths distance model (**c**). We generate a null model of expected over/under-citation rates, where the expected citation counts are the probabilities from the model that account for expected citation rates, and the observed citations counts are generated by pseudo-randomly citing based on those probabilities. For statistical inference, *p*-values are then calculated using this null distribution, and values that pass Holm-Bonferroni correction at *p* = 0.05 are annotated. (**d-f**) To visualize the intersections of race/ethnicity and gender in panels **a-c**, we plot the percentage point difference in over/under-citation, comparing men and women authors within each race (*gender*; grey), and comparing White authors to Asian, Black, and Hispanic authors (race; white) for three models: the random draws model (**d**), the paper characteristics model (**e**), and the shortest paths distance model (**f**).

### Temporal dynamics of co-authorship and its impact on over/under-citation

Prior work in other fields has suggested that homophily according to both gender and race/ethnicity exists in human relationships (41, 42); furthermore, gender homophily exists in scientific co-authorship (43–45). Should racial and ethnic homophily also exist in scientific co-authorship, either based on geographical or personal factors, it could produce biased perceptions of the overall racial and ethnic composition of a field. To assess this possibility, we built a time-evolving co-authorship network, thereby allowing us to evaluate whether authors of color are segregated into local co-authorship communities. In this network, edges exist between authors (including middle authors) who have co-authored a paper either that year or any year previously. To determine whether these true co-authorship patterns were explained by racial/ethic category, we constructed for each year 1000 partitions of authors into racial/ethnic categories consistent with the racial/ethnic category probabilities associated with each author. For example, if an author’s racial/ethnic category probabilities were 0.2 (Asian), 0.0 (Black), 0.1 (Hispanic), and 0.5 (White) then in most of the 1000 partitions they would be located in the White community, in some partitions they would be located in the Asian or Hispanic communities, and in none of the partitions would they be located in the Black community. To quantify the degree to which each of those probabilistic racial/ethnic partitions accounted for the co-authorship patterns, we used the *Q* statistic (see Methods).

As a null model that is naive to racial/ethnic category, we used a data-driven clustering method called InfoMap (46–48). The algorithm assigned authors to communities by maximizing the probability that authors in a single community are co-authors. Again, we quantified the degree to which this partition accounted for co-authorship patterns using the Q statistic (see Methods; (49)). We find that the data-driven partition exhibits its highest Q values from 1995-2000, thereafter decreasing (Figure 4). In contrast, the partition based on racial/ethnic category displayed a *Q*-value close to zero in 1995, which then increased dramatically over the subsequent two decades. This result suggests that, even as the clustering of authors in the network decreases in general, racial and ethnic segregation of authors of color is increasing over time.

### Co-authorship network model of over/under-citation

We next sought to test the hypothesis that authors tend to preferentially cite papers by authors that are close to them in the co-authorship network. For each paper, we therefore calculated (i) the distance (measured by the shortest path between two authors) between the first author and every other author, and (ii) the distance between the first author and every author that they cited. We performed the same two calculations for the last author. The average distance between a citing author and a cited author (mean=3.5) was significantly shorter (paired Student’s *t* = 418, −log10(p)>250, df=18,381) than the average distance between a citing author and all authors (mean=4.95). This result demonstrates that authors do tend to cite authors nearby in the co-authorship network.

Given this tendency, we sought to measure over/under-citation rates relative to the expectation that authors cite the papers of authors close to them in the co-authorship network. We built an expectation-rate model from the co-authorship network: the shortest path distance model, in which authors randomly cite based on the shortest path distance from the citing authors to the authors in the reference list (see Methods, Figure 5; for complementary results using a random walker model see Extended Data Figure 7 and 8f-h). For each paper, we generated a custom co-authorship network that only contained connections between authors who had co-authored a paper prior to the citing paper we are considering. We found very similar citation behavior to that we observed when we used the random draws and paper characteristics models to estimate the base rates. In the shortest path distance model (Figure 5b-d, Extended Data Figure 8c-e), White authors are still significantly over-cited, and papers led by authors of color are still significantly under-cited. Notably, the effect was driven more by White authors than by authors of color.

Next, we assessed whether authors’ tendency to cite papers closer or farther away within the co-authorship network (see Methods) is associated with the degree of imbalance among their citations. To assess this possibility, we first quantified the extent to which each citing team over-cited papers written by White first and last authors (WW papers), and then assessed whether authors’ average path length to citations was associated with that imbalance. On average, WW teams tended to cite other WW teams roughly 11% more than expected, mixed teams (WC and CW) cited WW teams roughly 6% more than expected, and CC teams cited WW teams roughly 1.5% less than expected. Among WW teams, the path length to citations was negatively associated with over-citation of WW papers (*β* = −0.98%, *t*_8958_ = −3.99, *p* < 0.0001), suggesting that more socially localized citation practices are associated with more over-citation of other WW teams. This association is a moderate one, however; WW teams with path lengths two standard deviations (SDs) above the overall average would still be expected to over-cite WW papers by roughly 8.8%. Conversely, path length to citations was *positively* associated with citation of WW papers among CC teams (*β* = 1.29%, *t*_2076_ = 2.24, *p* = 0.02). This positive association yields a slight over-citation of WW papers (by roughly 1%) among CC teams with path lengths two SDs above the overall average. Within mixed teams, path length to citations is not associated with citation behavior (*β* = 0.11%, *t*_4928_ = 0.332, *p* = 0.74). In general, these results demonstrate that citing further from oneself on the network is associated with more balanced citations. However, even WW teams with relatively distant citations tend to preferentially cite over WW teams. Extended Data Figure 9 visualizes the group-specific associations between path length and citation imbalance.

### The intersection of gender and race/ethnicity

In prior work, we examined the extent and drivers of gender imbalance in the reference lists of articles from these same five journals (33). Here, we seek to understand citation costs at the intersection of gender and race. In addition to a pair of racial/ethnic categories, each paper also has a pair of gender categories: man(first)-man(last), man-woman, woman-man, or woman-woman. Thus, instead of having 16 probabilities for each paper, we now have 64. We calculated the expected base-rates using the random draws model, the relevant characteristics model, and the shortest paths distance model. Consistently across all models, we observe a strong block diagonal structure indicating that men-led teams of all races are generally over-cited (see redder colors in the upper left square of Figure 6a-c), and woman-led teams of all races are generally under-cited (see darker blue in the lower right square of Figure 6a-c).

To quantify these observations, we considered papers with both first and last authors in a single racial/ethnic category and a single gender category; so, for example, we will use the term “women CC papers” to refer to papers whose first and last authors are women of color. In the random draws model (Figure 6a, Extended Data Figure 10a-d), we find a 31.06 (95%CI=7.73,60.15) percentage point gap between men CC papers and women CC papers. We find a 31.84 (95%CI=11.13,49.01) percentage point gap between White men papers and men CC papers. We find a 54.41 (95%CI=52.61,56.18) percentage point gap between White men papers and White women papers. We find an 8.47 (95%CI=-8.49,23.06) percentage point gap between White women papers and women CC papers. Finally, we find a 22.57 (95%CI=5.14,43.42) percentage point gap between men CC papers and White women papers.

Similar results are observed for both the paper characteristics model and the shortest paths distance model (Figure 6b-c, Extended Data Figures 11a-d, 12a-d). For the former, we find a 23.94 (95%CI=-0.64,45.09) percentage point gap between men CC papers and women CC papers. We find a 15.69 (95%CI=-2.24,27.28) percentage point gap between White men papers and men CC papers. We find a 29.06 (95%CI=27.08,31.0) percentage point gap between White men papers and White women papers. We find a 10.57 (95%CI=-4.19,19.69) percentage point gap between White women papers and women CC papers. Finally, we find a 13.37 (95%CI=-4.19,19.69) percentage point gap between men CC papers and White women papers.

In the shortest paths model, we find a 39.06 (95%CI=7.8,78.48) percentage point gap between men CC papers are and women CC papers. We find a 33.72 (95%CI=10.06,58.49) percentage point gap between White men papers and men CC papers. We find a 66.69 (95%CI=65.02,68.38) percentage points gap between White men papers and White women papers. We find a 6.06 (95%CI=-14.3,22.56) percentage points gap between White women papers and women CC papers. Finally, we find a 32.97 (95%CI=7.99,56.79) percentage point gap between men CC papers and White women papers.

In the supplement, we show that the results also hold in the random walker model (Extended Data Figure 7 and 8f-h). Finally, we separately measure the relative impacts of gender and race/ethnicity on over/under-citation (Figure 6d-f, Extended Data Figures 10e-h, 11e-h, 12e-h).

## Discussion

Academia and industry harbor racial and ethnic discrimination just as ubiquitously as other areas of our society (1, 2, 4). Discrimination is a common thread in the fabric of scientific culture. It arises from both implicit bias and explicit bias. In the former, subconsciously harbored discriminatory attitudes can manifest in response to someone’s person or in response to someone’s name, which is used to infer features of the person (50). Bias against racial and ethnic minorities leads to marked disparities, which exist in promotion, retention, grant funding, awards, and publications (5–10). Such disparities are often viewed as the responsibility of people in positions of power (e.g., journal editors, society presidents, award committee members, conference organizers). Yet, many disparities are perpetuated by researchers at all levels. Citation practices are a particularly good example. As the currency in academia (28), citations strongly influence career advancement, and as a record of science, citations reflect the value we place on scientific questions and their questioners.

Here we set out to assess the presence and extent of racial and ethnic imbalance in citations in the reference lists of five top neuroscience journals from 1995 to 2019. Using previously developed race prediction models (38, 39), we assigned each author’s name a probability of being held by a person of a given racial/ethnic category. We found that papers written by authors of color are under-cited relative to the proportion of such papers in the field, and that the under-citation is largely being driven by White authors. Notably, undercitation of authors of color is increasing with time, despite the growing diversity of the academy. The most imbalanced citers are those who cite authors nearby on the co-authorship network, which itself is becoming increasingly segregated by race over the last 25 years. Future work could expand the assessment of social factors influencing citation behavior by examining author’s academic heritage (e.g., graduate and postdoctoral advisors) using for example *neurotree*.

Our findings are markedly consonant with prior work assessing gender (rather than racial or ethnic) imbalance in references lists across several fields including neuroscience (33), international relations (30), astronomy (31), and political science (29, 32). In those studies, the gender minority (women) were under-cited (29–33), and that under-citation was largely driven by the gender majority (men) (29, 32, 33); just as here, we find that the racial/ethnic minority (authors of color) are under-cited, and that under-citation is largely driven by the racial/ethnic majority (White authors). Moreover, in neuroscience the under-citation of the minority gender (women) is increasing with time, particularly in the reference lists of the gender majority (men) (33); just as here, we find that the under-citation of the racial/ethnic minority (authors of color) is increasing with time, more strongly in the reference lists of the racial/ethnic majority (White authors). The consistent effects of gender and race/ethnicity on citations suggests the possibility of shared biases to peoples that are minoritized along any of several potential dimensions of difference. Future work could investigate this possibility by expanding to other dimensions of difference.

Importantly, we observe that the penalty of undercitation in our scientific culture is being paid more by women than by men. When considering the degree of over/under-citation of first and last authors, of the four racial/ethnic categories and the two genders studied here, we find that the dominant mode of variation is along the line of gender, whereas the secondary mode of variation is along the line of race/ethnicity. White men are most over-cited. Men of color are less over-cited than White men, although they are not under-cited on average. Women of all racial/ethnic categories are under-cited; Black women are the most undercited. This set of findings may guide ongoing and future efforts to mitigate disparity in the academy; it suggests that we must engage much more consciously and conscientiously with the work of people of color, and even more with the work of women scientists of any racial/ethnic background, and even more with the work of Black women.

### Tracking race and ethnicity

The precise definitions of race and ethnicity remain a topic of much debate. While race is commonly taken to signify shared ancestry and physical appearance, such as facial features or skin color, and ethnicity is commonly taken to signify shared ancestry and cultural features, such as language or religion, most scholars agree that both are dynamic labels that are societally generated, embodied, policed, and revised (51). Race and ethnicity, then, become a set of social, political, and/or cultural groupings of people. Although some scholars argue there is a biological basis to race, most agree that there is no biological basis either to race or to the social hierarchies of races (52), and that in fact the purported biological basis of race is itself socially constructed alongside the hierarchies it is supposed to justify (53). The specific architectures of those social constructions, as the result of economic and political processes, are often termed “racial formations” (54). Race and ethnicity are not only socially crafted, however; they are also subjectively felt (55). Race and ethnicity are phenomenologically real; they are things people regularly experience. One of the ways race and ethnicity are experienced is through biases against visible or aural differences to which racial and ethnic meanings have been assigned. Dress, posture, language, dialect, accent, facial features, skin color, names, and even ambitions are taken as codes or ciphers of race and ethnicity. That is to say, they are racialized or ethnicized.

In this study, we focus on the racialization and ethnicization of names. For our purposes, “race/ethnicity” refers to the probability of a name being assigned to someone who identifies or is identified as belonging to a specific racial/ethnic category. Probabilities of being “Asian,” “Black,” “Hispanic,” and “White” are assigned to each author based on the prevalence of their first/last name bicharacters in people identifying or identified as “Asian,” “Black,” “Hispanic,” and “White” in the name datasets. The actual racial or ethnic identification of the author is not identified. Given the limitations of probabilistic analyses, the authors may in fact have a racial or ethnic identity different from the one we have assigned and/or another racial/ethnic identity in addition to the one we have assigned. In some cases, citers will know the racial or ethnic identification of the authors they cite. In many cases, they will not know, but rather infer the racial/ethnic category of the authors they cite. Instances of both known and inferred racial/ethnic identification have the potential to incite either explicit or implicit bias in citing authors. Our probabilistic analysis by racialized and ethnicized name, therefore, functions to non-trivially capture bias arising due to both known and inferred racial/ethnic identification in citation practices.

Bias, which refers to a distortion of judicial treatment, can be understood along two vectors: from structural to individual, and from implicit to explicit. Structural bias refers to discriminatory values, practices, and mechanisms that function at the intergroup level in the domain of social institutions. Individual bias refers to discriminatory values, practices, and behaviors that function at the interpersonal level in the domain of social interaction. Both structural and individual bias may be either unconscious or conscious, furtive or overt. Racial and ethnic bias, in particular, refers to oppressive, discriminatory, or prejudicial values and practices that supervene on a person’s known or presumed racial or ethnic identification. Manifestations of structural bias with respect to race and ethnicity include the over-policing and over-incarceration of communities of color (56, 57); in academia, they also include hiring and retention practices in which racial struggles remain unaccounted for while diversity service work (with which faculty of color are overburdened) routinely counts against reappointment, tenure, and promotion (19, 58). Manifestations of individual bias with respect to racialized or ethnicized names may include not hiring someone with a non-Anglophone name (17, 18). In academic contexts, both explicit and implicit racial/ethnic bias may result in passing over a person of color for a hire, promotion, grant award, or collaborative opportunity, or considering their work as likely having less impact, value, or scholarly rigor and excellence. Our study considers the under-citation of authors of color as an indicator of racial and ethnic bias in neuroscience.

Racial and ethnic bias cannot be understood in abstraction from biases of gender, disability, sexuality, and class. By intersecting, these biases complexify and compound one another. Women of color have historically led the theorization of intersectionality (59–63). Kimberlé Crenshaw famously argues that discrimination can come from multiple directions at once and intersect to devastating effect. As such, the “intersectional experience” of a specifically racialized, ethnicized, gendered, and classed person “is greater than the sum” of each axis (64). Recent work focusing on working class women of color faculty illustrates this compound effect. Despite the specificities of their differences as women (of a variety of gender presentations), people of color (of a variety of racial or ethnic identities), and working class, women of color faculty with working class backgrounds are often hyper-marginalized and disrespected. They are “presumed incompetent” in the classroom, in their labs, as writers and speakers, as organizers and leaders (19, 65). Our findings that women of color authors are significantly more undercited than White men and men of color, and are slightly more under-cited than White women, confirms the compound effects at the intersection of gender and racial/ethnic bias. We hypothesize that future work accounting for additional axes of bias would see a similar compounding of under-citation.

### Methodological considerations and limitations

We wish to acknowledge several important limitations and complexities of the methods we employ. First, the Census categories themselves are historico-political projects long entangled with colonization, slavery, miscegenation laws, blood quantum laws, and assimilation pressures for Indigenous peoples and people of color (66, 67). This means that people are counted and count themselves as belonging to racial/ethnic categories to which they may or may not belong. Second, the most recent Census data was collected 10 years ago, and much has changed in the demographics of the United States since. Third, the Census and Florida models drawn from the Census data and Florida Voter Registration data do not calculate data for American Indian/Native Alaskan or mixed-race people. This means that Indigenous and mixed-race authors are not explicitly modeled in the source data. However, in our analysis, if an author has, for example, a first name that is common to one racial/ethnic category, and a last name that is common to another racial/ethnic category, this fact will be reflected by that author having high probabilities for both racial/ethnic categories. Fourth, we acknowledge that the racial/ethnic categories studied here are coarse and future work would do well to examine finer categories, paying particular attention to people groups with diverse and ambiguous racial/ethnic histories (68–70). Fifth, our approach does not explicitly measure how citation practices might vary among people of different races and ethnicities. We look forward to future work that extends the present analysis to Indigenous and mixed-race authors, as well as better accounts for authors of color who may face differential biases due to the ambiguous racialization or ethnicization of their names.

### Looking to the future

When faced with clear evidence of inequality, we must each determine what we will do in response. In many areas of academic disparity, the conversation revolves around building better pipelines to ensure a diverse faculty. Yet, here we find that the growing diversity of bodies in academia is accompanied by a growing segregation of minds. Moreover, prior work clearly demonstrates that greater representation cannot be assumed to reduce discrimination (21). Hence, the efforts to physically diversify the bodies we place in university offices must be complemented by efforts to mentally diversify the ideas and idea-makers that we engage with, cite, and place before our students in colloquia, syllabi, hallway portraiture, *et cetera* (71, 72). Diverse ideas are the life blood of science; they are the steps we take to optimally forage in the vast space of the unknown. By not citing questioners based on their gender or race/ethnicity, we limit the questions being asked thereby fundamentally handicapping the progress of science.

To hasten the scientifically vigorous future that can only arise in an equitable culture, we must embrace personal responsibility. For those not paying the highest costs to engage in a scientific career, it is imperative to define what allyship means (22), and educate our faculty and trainees accordingly (23, 24). It will not suffice to mentor gender and racial/ethnic minority scholars to meet the challenges they face (25); rather non-minority scholars must acknowledge that they create those challenges and perpetuate them.

All scholars, but particularly scholars who are not minorities along the dimensions of race/ethnicity or gender, can choose to cite equitably, or even to cite in a manner that actively counteracts the long histories of discrimination that these people have and continue to face. For those readers interested in inquiring about their own citation practices using the tools laid out here, see freely available software executable from any internet browser (35) as well as open source Python scripts (73). For those more broadly seeking actionable recommendations for hastening an equitable future, see Refs. (34, 74). Fundamentally, citations are “a reproductive technology, a way of reproducing the world around certain bodies” (75). Will we reproduce what we have inherited? Or will we join together to transform the practice of citation into a practice of conscientious engagement (76), thereby paving the way for a new kind of scientific culture in which each voice is heard equally?

## Acknowledgments

We are grateful to the many mentors, colleagues, and mentees who have cultivated our own awareness of disparity over the years and our commitment to its mitigation. DSB would like to acknowledge support from the John D. and Catherine T. MacArthur Foundation and the National Science Foundation (CAREER PHY-1554488 and IIS-1926757). DSB and PZ acknowledge additional support from the Center for Curiosity. The content is solely the responsibility of the authors and does not necessarily represent the official views of any of the funding agencies.

## Methods

### Data collection

For this study, we drew data from the Thomson Reuters’ Web of Science (WoS) database which indexes neuroscience journals according to the Science Citation Index Expanded. To select journals, we used the Eigenfactor scores, which give a count of incoming citations weighted by the impact of the citing journal. The Eigenfactor score roughly mimics the classic version of Google page rank, and attempts to characterize the influence that a journal has within its field (37). We chose the five neuroscience journals with the highest Eigenfactor scores: *Brain, Journal of Neuroscience, Nature Neuroscience, NeuroImage*, and *Neuron*. For each journal, we downloaded all articles published between 1995 and 2019 that were classified as either empirical articles, review articles, or proceedings papers. The data included papers’ author names, reference lists, publication dates, and Digital Object Identifiers (DOIs). To study citation structure among papers, we matched the DOIs contained within a reference list to the DOIs of papers included in the dataset. We chose 1995 as the starting year because *NeuroImage* was founded in 94-95 and *Nature Neuroscience* was founded in 98; the mid-1990s is thus the point at which the newer target journals came into existence. The mid-1990s also marks a reasonable inflection point for publication output across the journals.

Although authors’ last names were included for all papers, authors’ first names were only regularly included for papers published after 2006. For all papers published in or before 2006, we searched for author first names using Cross-ref’s API. When first names were not available on Crossref, we searched for them on the journals’ webpage for the given article. To minimize the number of papers for which we only had access to authors’ initials, to remove self-citations, and to develop a co-authorship network, we implemented a name disambiguation algorithm.

### Author name disambiguation

To minimize missing data, allow for name race/ethnicity and gender assignment, and allow for author matching across papers, we implemented an algorithm to disambiguate authors for whom different versions of their given name or initials were present across papers. We began by separating first and last names according to the method used by the given source (e.g., WoS typically used “last, first; last, first”). We then identified cases in which only initials were available after the previously described searching steps by marking authors for whom the first name entry contained only uppercase letters (as we found that many initials-only entries did not contain periods).

For each case, we collected all other entries that contained the same first/middle initials and the same last name. If only one unique first/middle name matched the initials of the given entry, or if distinct matches were all variants of the same name, we assigned that name to the initials. If there were multiple names in the dataset that fit the initial/last name combination of the given entry, then we did not assign a name to the initials. For example, if an entry listed an author as R. J. Dolan, and we found matches under Ray J. Dolan and Raymond J. Dolan, we would replace the R. J. Dolan entry with the more common completed variant. If, instead, we found matches under Ray J. Dolan and Rebecca J. Dolan, we would not assign a name to the original R. J. Dolan entry.

Next we matched different name variants for the sake of tracking individual authors across their papers. To find and connect variants, we searched for instances of author entries with matching last names and either the same first name or first names that were listed as being commonly used nicknames according to the Secure Open Enterprise Master Patient Index (77). If there were no matches that fit that description, the name was retained. If there was one match that occurred more commonly, the less common variant was changed to the more common variant. If there were multiple matches that did not have any conflicting initials (some having a middle initial and others not having one was not considered conflicting), then less common variants were changed to the more common variant. If there were multiple matches that did have conflicting initials (e.g., Ray Dolan being matched to both Raymond S. Dolan and Raymond J. Dolan), then the target name was not changed.

There are three primary ways that incorrect author disambiguation could impact the results presented in this study. First, inability to link initials to an author’s first name would yield missing data for papers that only included the author’s initials. These papers would then not be included in the analyses as either *cited* or *citing* papers. Second, inability to link two versions of an author’s name (e.g., Ray Dolan and Raymond Dolan) would lead to the inclusion of some self-citations into the author’s analyzed reference lists. This inclusion could lead to slightly inflated rates of authors citing other authors of the same race/ethnicity or same gender, though sensitivity analyses suggest that this effect is essentially nonexistent in the present data (33). Third, incorrectly linking Author A to Author B would lead to the unnecessary removal of some citations (i.e., any of Author B’s references to Author A’s work would be removed as self-citations). Though this is likely a rare occurrence, its presence would lead to slightly decreased rates of authors citing other authors of the same race or same gender.

### Author race determination

We utilize two datasets to assign a racial/ethnic category to an author’s name; we present one in the main text and the other in the Extended Data. In the main text, we utilize a dataset collected and built by others (38), where hidden Markov models and decision trees were used to classify first and last names into racial/ethnic groups: Asian/Pacific Islander, non-Hispanic Black, Hispanic, and non-Hispanic White. Note that we use the term Hispanic (referring to native speakers of Spanish or those with Spanish-speaking ancestry) rather than Latinx (referring to those of Latin American origin or ancestry) because the prediction algorithm evaluates the linguistic content of names; we acknowledge that some of the authors in our database may identify as Latinx. In the Extended Data, we use data from the Census Bureau, which provides the primary racial/ethnic category associated with all last names occurring 100 or more times for the 2000 and 2010 census.

We used the open source python package, ethnicolr (39), to model the predicted racial/ethnic category for specific names. The model steps can be summarized as follows. (1) Concatenate last name (and first name in the full name Florida Voter Registration model) and capitalize the first character of all words. (2) Split the name into two character chunks (bi-chars); for example “Smith” would be split into “Sm”, “mi”, “it”, and “th”. (3) Remove infrequent bi-chars defined as those occurring less than 3 times in the data, and remove very frequent bi-chars defined as those occurring in more than 30% of the data. (4) Sort by frequency and build up the words list (bi-chars). (5) Build X as the index of bi-chars in the words list. (6) Pad the sequences in X so that they are the same size: 20 for the last name only model and 25 for the full name model. (7) Split into train and test: 80/20 and perform out of sample validation. (8) Train the model with long short term memory networks (LSTMs), which are recurrent neural networks capable of learning long-term dependencies.

The Florida and Census models differ in their accuracy and specificity (39). Across cross-validation folds, the average precision, recall, and f1-support scores for the Florida model were 0.83, 0.84, 0.83, respectively (39). No large biases were detected in the confusion matrix, which encodes the probability of predicting a person’s race to be X when their true race is X, as well as the probability of predicting a person’s race to be Y when their true race is X. In the Census model, the average precision, recall, and f1-support scores were 0.78, 0.80, 0.76, respectively (39). The Census model has the marked deficiency of frequently misclassifying Black and Hispanic people as White (39) (Figure 13).

The ethnicolor algorithm (39) can be used to make binary predictions but also provides probabilities that an author belongs to each racial/ethnic category. The probabilities are particularly useful for our study. Consider the last name “Smith” which is racially and ethnically ambiguous, particularly in the Census model without first name information. The model’s probabilities for the name “Smith” are 73% White, 25% Black, 1% Hispanic, and <1% Asian. We can use all four probabilities to estimate how citers probabilistically assign racial/ethnic categories to names, either implicitly or explicitly, while reading and citing papers. This probabilistic assessment is also useful when evaluating authors whose names are particularly complex, such as those whose first name is common in one race/ethnicity and whose last name is common in another race/ethnicity. Note that imperfections in the algorithm’s predictions will break the links between citation behavior and author race/ethnicity, and therefore any incorrect estimation in the present data likely biases the results towards the null model.

### Author gender determination

As described in our earlier study (33), for authors with available first names, gender was assigned to first names using the ‘gender’ package in R (78) with the Social Security Administration (SSA) baby name dataset. For names that were not included in the SSA dataset, gender was assigned using Gender API (gender-api.com), a paid service that supports roughly 800,000 unique first names across 177 countries. We assigned ‘man’(‘woman’) to each author if their name had a probability greater than or equal to 0.70 of belonging to someone labeled as ‘man’(‘woman’) according to a given source (32). In the SSA dataset, man/woman labels correspond to the sex assigned to children at birth; in the Gender API dataset, man/woman labels correspond to a combination of sex assigned to children at birth and genders detected in social media profiles. In our prior study, we selected a random sample of 200 authors, and found the accuracy of these automated gender assignments to be 0.96 (33). Note that imperfections in the gender prediction algorithm will break the links between citation behavior and author gender, and therefore any incorrect estimation in the present data likely biases the results towards the null model.

Here, gender could be assigned to both the first and last author of 88% of the papers in the dataset (n=56,144). To assimilate these categories into our probabilistic model of expected citation rates, we set the model’s gender weights to 0 for woman and 1 for man. Of the 12% of papers with missing data, 7% were missing because either the first- or last-author’s name had uncertain gender, and 5% were missing because either the first- or last-author’s name was not available. For these cases, we used gender weights equal to the base rates of author gender categories for papers published that year; for example the weight for a man first author is the proportion of papers that year that had a man first author.

Given the limitations of probabilistic analyses, the authors may in fact have a sex or gender different from the one we have assigned and/or be intersex, transgender, or nonbinary (79, 80). In some cases, citers will know the sex and/or gender of the authors they cite. In many cases, they will not know but rather infer the gender of the authors they cite. Instances of both known and inferred gender have the potential to incite either explicit or implicit bias in citing authors (16, 18, 81). Our probabilistic analysis by gendered name therefore functions to nontrivially capture bias in citation practices arising due to both known and inferred gender.

### Co-authorship network models and null model

In the main text, we evaluated the role of co-authorship networks in citation practices by defining a shortest paths distance model. Here, for each citing paper, we calculated the distances between (i) the first and last author of the citing paper, and (ii) the first and last authors cited in that paper’s reference list (Fig. 5 and Extended Data Fig. 8a). For each citing paper, this process creates a distribution of the distances between the two citing authors and the cited authors. To create a pool of similar papers that the citing authors could have cited, we randomly chose 1000 *potential* cited authors such that the distribution of distances was maintained. From each *potentially* cited author, we randomly chose one paper that was published prior to the true citing paper. The model thus builds base rates from the expectation that citing authors randomly cite papers of other authors that are the same distance away from them in the co-authorship network as those they did cite.

In the Extended Data, we complement the shortest paths model with a random walker model. Here, for the first (and then last) author of each paper, we start by finding the longest distance between that author and all other authors in the network. This quantity measures the distance that a random walker would need to travel to reach the author that is the farthest away in the co-authorship network. We then create a random walker, and program it to continue walking until it has visited this precise number of nodes. This process ensures the *possibility* that the random walker can reach any author in the network from the citing author. We start the random walker at the first (and then last) citing author and allow it to walk through the co-authorship network (Fig. 7a and Extended Data Fig. 8b). Once the preset number of nodes visited is reached, we randomly choose one paper from the author that the random walker reached in its final step. We performed the same approach for 1000 random walkers for the first (and then last) author, totalling 2000 walks per citing paper. The model thus builds base rates from the expectation that citing authors randomly cite papers of other authors that are close to them in the co-authorship network.

To perform statistical inference, we generated a null distribution of over/under-citation. For each citing paper, we generated null expected citation counts defined as the probabilities from the shortest paths distance model (main text) or random walker model (Extended Data). Then, we generated null observed citation counts by pseudo-randomly citing based on those probabilities. This null model thus reflects the possible distribution of over/under-citation values present while citing based on the network structure of co-authorship.

### Statistical modeling

For many analyses conducted in this study, we used a paper characteristics model that built expected citation rates by racial/ethnic category, or by both racial/ethnic and gender categories. When building this model solely for racial/ethnic categories, we fit a ridgeregression model (sklearn: RidgeCV) on the multinomial outcome of the 16 different combinations of author race (4 racial/ethnic categories, 2 author positions), in which the model’s features were 1) month and year of publication, 2) combined number of publications by the first and last authors, 3) number of total authors on the paper, 4) the journal in which it was published, 5) whether it was a review paper, and 6) the continent (i.e., location) of the corresponding author. Note that the latter is important as prior work indicates that differences between the country of the author and the country of an evaluator (e.g., journal editor or in our case citer) can impact the treatment of the publication (e.g., accept/reject decision or in our case citation count) (82). Similarly, when building the model for both racial/ethnic and gender categories, we fit a ridge-regression model on the multinomial outcome of the 64 different combinations of author gender and race/ethnicity (4 racial/ethnic categories, 2 genders, 2 author positions), in which the model’s features were the same (1-6) listed above.

For both models, the proper L2 regularization was found by performing cross-validation, with values ranging from 1 × 10^-5^ to 100. When the model is then applied to each paper, it yields a set of probabilities that the paper belongs to the 64 racial/ethnic and gender pair categories (for the intersectionality analyses), or to the 16 racial/ethnic pair categories (for the remaining analyses), respectively. Importantly, this process does not predict the number of citations given to individual papers. Instead, it facilitates the calculation of the rates at which different racial/ethnic and gender categories would be expected to appear in reference lists if author race and gender were independent of citation rates, conditional on the other characteristics in the model.

Throughout the study, we present estimates with confidence intervals that were calculated by bootstrapping citing papers. In each bootstrap, we take a random sample of citing papers with replacement. In contrast to bootstrapping individual instances of citations, this method maintains the dependence structure of the clusters of cited articles within citing articles.

### Hypothesis testing

Here, we describe the formal statistical analysis that we used to address the five distinct hypotheses. All hypotheses were tested for the set of articles published between 2009 and 2019. We considered reference lists from the past 10 years to ensure that estimates of over/under-citation reflected current behavior, were not a result of aggregating over disparate eras of neuroscience research, and contained enough prior citable papers to represent meaningful and stable measures of behavior.

#### Hypothesis 1: The overall citation percentage of C∪C papers will be lower than expected given papers’ relevant characteristics

To test the hypothesis that the overall citation percentage of papers by authors of color will be lower than expected given papers’ relevant characteristics, we first estimated the expected number of citations given to each racial/ethnic category. We obtained this expectation by calculating the sum of the model-estimated probabilities for all papers contained within the reference lists of citing papers. This sum, therefore, reflects the expected number of citations given to each racial/ethnic category if author race were conditionally independent of citation behavior, given the paper’s characteristics. These expectations form the model’s base rates.

To calculate the observed percentage of citations given to each racial/ethnic category, we took the sum of the probabilities of each author racial/ethnic category for cited papers in the papers’ reference lists. Notably, performing the summation over all citations results in the upweighting of articles with many citations, and the downweighting of articles with few citations. This approach helps to improve the stability of the estimates (33). We then converted the model’s base rates and the observed rates into percentages. Now, the observed values can be compared to the base-rate values by calculating the percent difference from expectation for each author racial/ethnic category. For example, for White-White (WW) papers, this percent change in citation would be defined as,

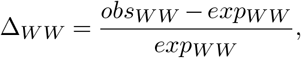

where *obs_WW_* is the number of citations given to WW papers between 2009 and 2019, and *exp_WW_* is the expected number of citations given to WW papers between 2009 and 2019.

#### Hypothesis 2: Under-citation of papers led by authors of color will occur to a greater extent within White-led reference lists

To test the hypothesis that the under-citation of papers led by authors of color will occur to a greater extent within the reference lists of White-led teams, we used very similar metrics to those described in the previous section. The primary difference is that instead of calculating the observed and expected citations by summing over the citations within all reference lists between 2009 and 2019, here we performed those summations separately for reference lists in papers with White first and last authors (WW papers) and papers with at least one author of color as the first or last author (CUC papers). For example, to estimate the over/under-citation of C∪C papers within the reference lists of WW papers, we define

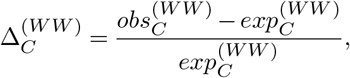

where 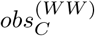 is the total number of citations given to C∪C papers within WW reference lists, and 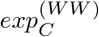 is the expected number of citations given to C∪C papers within WW reference lists.

#### Hypothesis 3: Under-citation of C∪C papers will be decreasing over time, but at a slower rate within White-led reference lists

To evaluate change in citation balance over time, we calculated the over-citation of, for example, White authors over time using the simple measure of the absolute difference between the observed proportion of WW papers cited and the expected proportion of WW papers cited. This measure of change is given by,

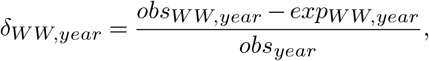

where *obs_year_* is the total number of citations within a given year, *obs_WW,year_* is the number of citations given to WW papers in a specific year, and *exp_WW,year_* is the expected number of citations given to WW papers in a specific year.

#### Hypothesis 4: The under-citation of C∪C papers by White citers will be partly explained by White authors citing authors nearby on the co-authorship network

We operationalized this hypothesis by determining whether authors’ tendency to cite papers closer or farther away within the co-authorship network is associated with the degree of imbalance among their citations. We quantified the extent to which each citing team over-cited papers written by White first and last authors (WW papers), and then assessed whether authors’ average path length to citations was associated with that imbalance. The method allows us to determine whether more socially localized citation practices are associated with more over-citation of other WW teams. Given that the individual estimates of each paper’s over/under-citation are noisy, we used the distances and citation rates obtained from each of the 10,000 bootstrapped samples of the paper characteristics model.

#### Hypothesis 5: Under-citation of C∪C papers will be greater for women of color than for men of color

In our final analysis related to intersectionality, we use the same approach as described in *Hypothesis 1* to calculate over/under-citation for author racial/ethnic and gender categories. Specifically, we calculate *exp_WW_* and *obs_WW_* separately for all 64 combinations of author genders and race/ethnicity. Intuitively, these calculations provide the expected number of citations given to each author racial/ethnic and gender category, as well as the observed percentage of citations given to each gender and racial/ethnic category. Next, we generate a 95% confidence interval for each comparison (e.g., White-men papers *versus* White-women papers) by measuring the differences between bootstrap means for each racial/ethnic and gender category we tested.

### Time-evolving segregation of co-authorship network

In our assessment of the time-evolving segregation of authors of color in the co-authorship network, we compared a data-driven partition to a partition based on race/ethnicity alone. In both cases, the quality of the partition was quantified by the modularity quality index:

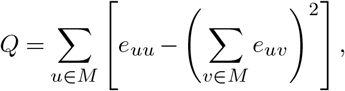

where the network is partitioned into a set of non-overlapping modules *M*, and *e_uv_* is the proportion of all links that connect nodes in module *u* with nodes in module *v* (49). The quantity *Q* allowed us to assess racial/ethnic segregation in the coauthorship network as a function of time.

## Citation Diversity Statement

Recent work in several fields of science has identified a bias in citation practices such that papers from women and other minority scholars are under-cited relative to the number of such papers in the field (29–33). Here we sought to proactively consider choosing references that reflect the diversity of the field in thought, form of contribution, gender, race, ethnicity, and other factors. First, we obtained the predicted gender of the first and last author of each reference by using databases that store the probability of a first name being carried by a woman (33, 35). By this measure (and excluding self-citations to the first and last authors of our current paper), our references contain 39.65% woman(first)/woman(last), 11.99% man/woman, 17.99% woman/man, and 30.37% man/man. This method is limited in that a) names, pronouns, and social media profiles used to construct the databases may not, in every case, be indicative of gender identity and b) it cannot account for intersex, non-binary, or transgender people. Second, we obtained the predicted racial/ethnic category of the first and last author of each reference by using databases that store the probability of a first and last name being carried by an author of color (38, 39). By this measure (and excluding self-citations), our references contain 12.55% author of color (first)/author of color(last), 17.55% white author/author of color, 15.78% author of color/white author, and 54.12% white author/white author. This method is limited in that a) names and reported race/ethnicity used to make the predictions may not be indicative of racial/ethnic identity, and b) it cannot account for Indigenous and mixed-race authors, or those who may face differential biases due to the ambiguous racialization or ethnicization of their names. We look forward to future work that could help us to better understand how to support equitable practices in science.

## Supplementary Information

**Extended Data Figure 1.**
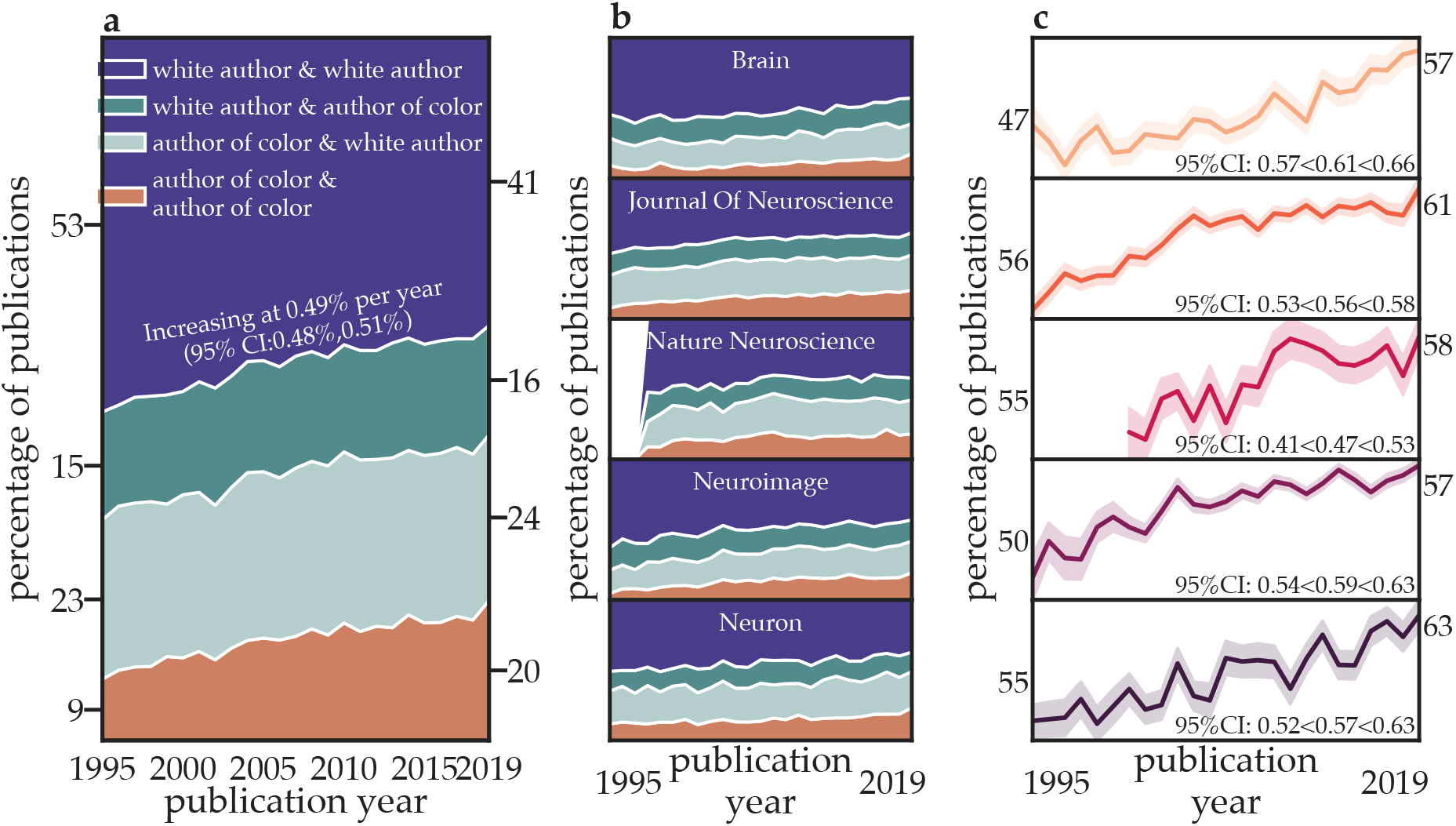
Racial and ethnic diversity is increasing within top neuroscience journals (Florida model; data separated by racial/ethnic category). **a** The percentage of papers published by authors of distinct racial/ethnic categories across the five journals studied from 1995 to 2019. **b** The percentage of papers published by authors of distinct racial/ethnic categories in each journal separately from 1995 to 2019. **c** The percentage of papers with a first and/or last author of color for each journal, for each year. Confidence intervals for the change in percent of papers with a first or last author of color (C∪C) for each year and for each journal were generated via bootstrapping (*n*=1000). Note: The journal *Nature Neuroscience* was not established until 1998.

**Extended Data Figure 2.**
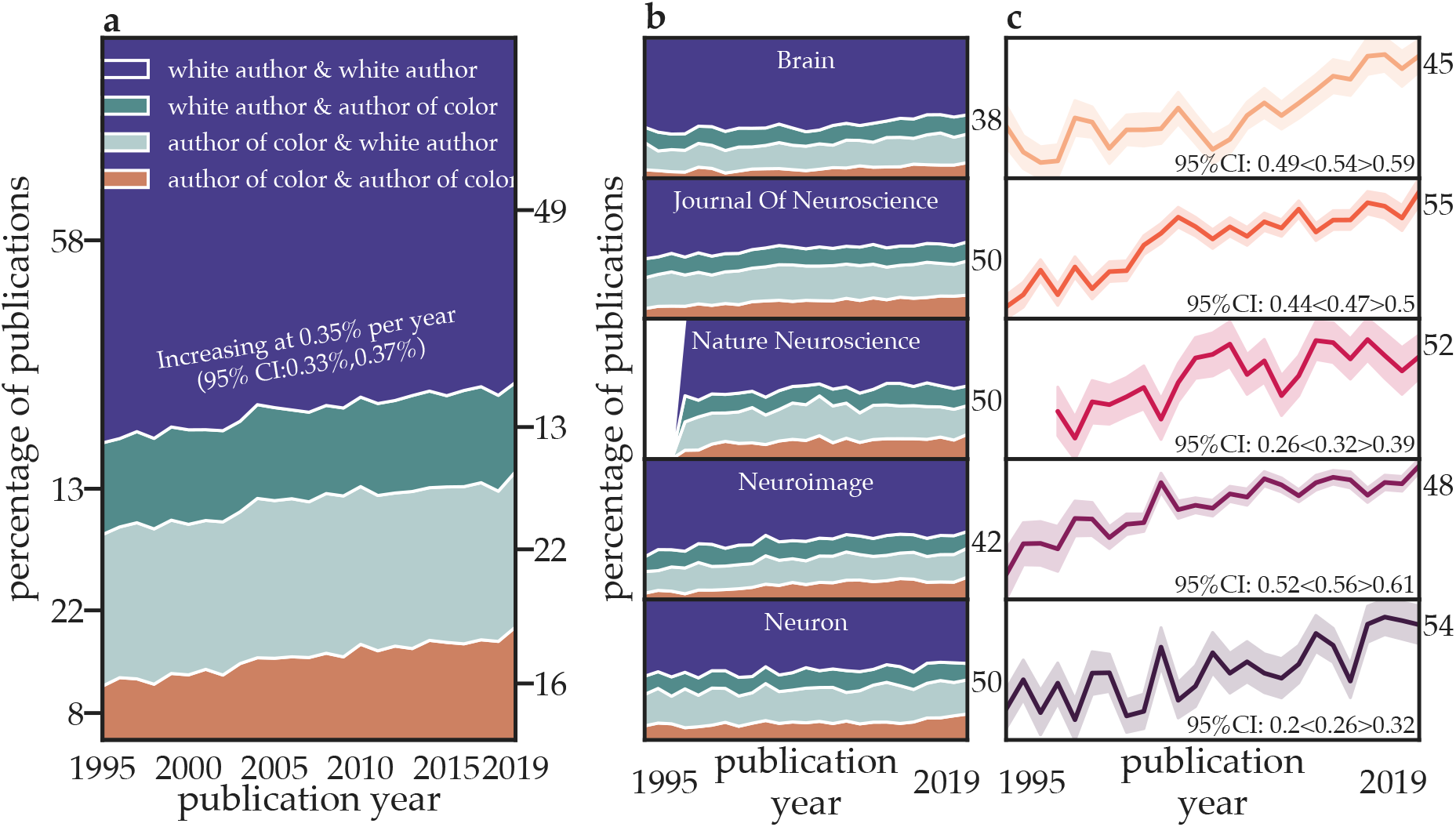
Racial and ethnic diversity is increasing within top neuroscience journals (Census model; data separated by racial/ethnic category). **a** The percentage of papers published by authors of distinct racial/ethnic categories across the five journals studied from 1995 to 2019. **b** The percentage of papers published by authors of distinct racial/ethnic categories in each journal separately from 1995 to 2019. **c** The percentage of papers with a first and/or last author of color for each journal, for each year. Confidence intervals for the change in percent of papers with a first or last author of color (C∪C) for each year and for each journal were generated via bootstrapping (n=1000). Note: The journal *Nature Neuroscience* was not established until 1998.

**Extended Data Figure 3.**
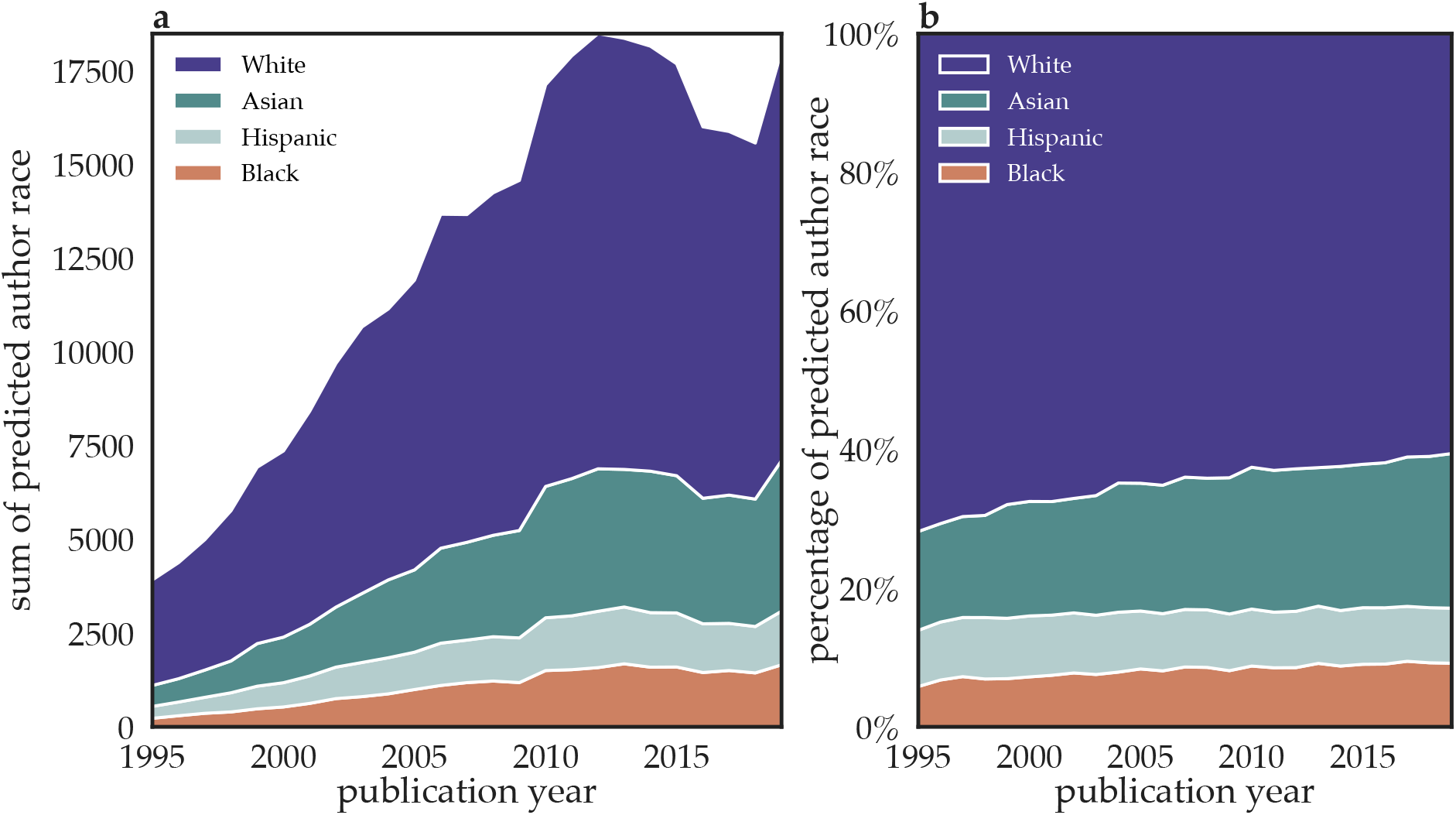
The authors publishing in top neuroscience journals are increasingly diverse in terms of race and ethnicity (Florida model). **a** The sum of probabilities of distinct racial/ethnic categories of the first and last authors of papers published in the top five neuroscience journals studied from 1995 to 2019. **b** The percentage of the total author pool that is comprised of authors in the four racial and ethnic categories studied.

**Extended Data Figure 4.**
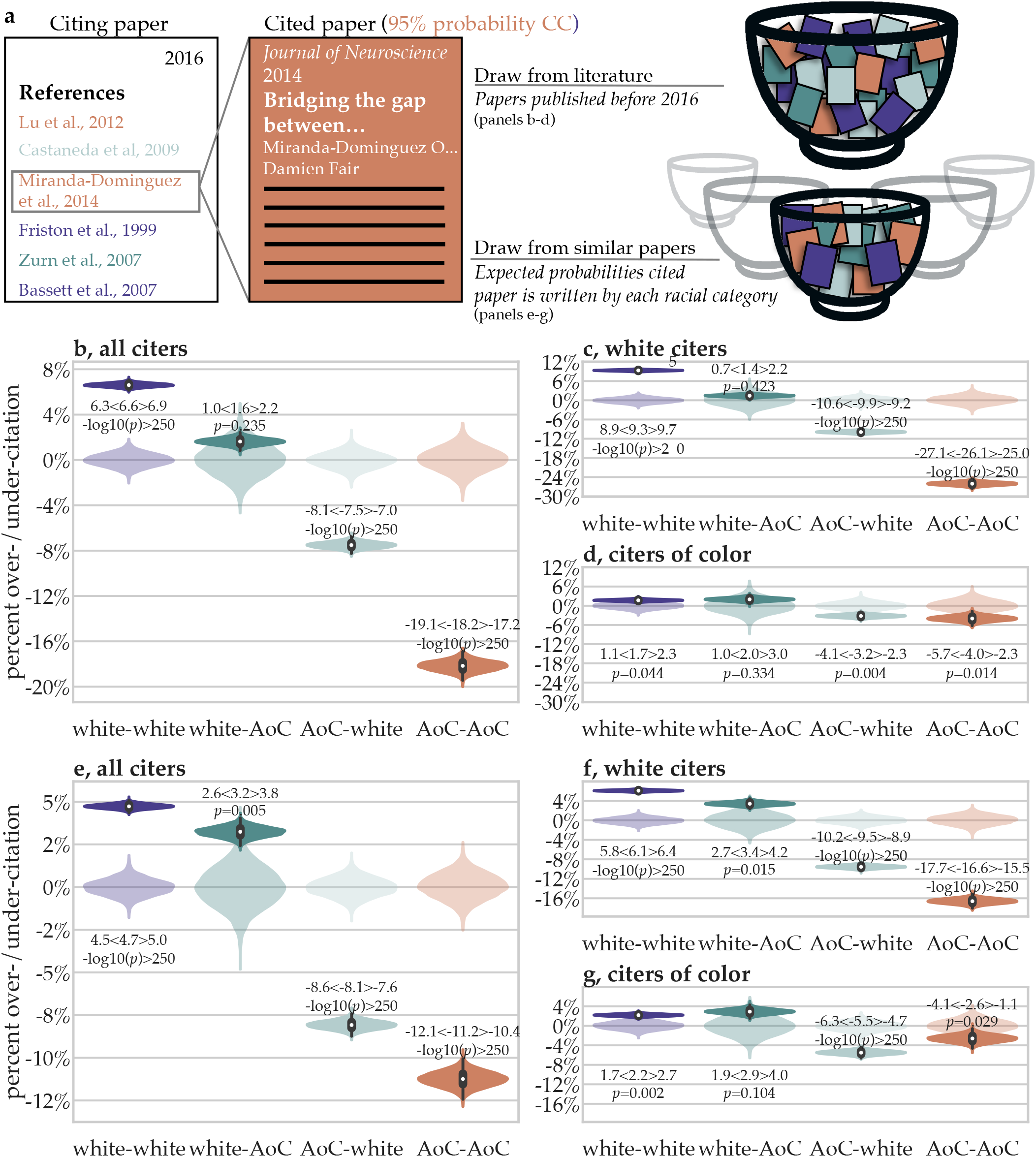
Models and results for the over/under-citation of papers based on predicted author race/ethnicity (Census Model). **a** In the random draws model, racial/ethnic category proportions in reference lists are compared to the overall racial/ethnic category proportions of the existing literature. In the relevant characteristics model, author category proportions in reference lists are compared to author category proportions of similar articles by 1) year of publication, 2) journal of publication, 3) number of authors, 4) research article or review, 5) first and last author seniority, and 6) the location of the authors’ institution. **b** The over/under-citation of different author categories compared to their expected proportions under the random draws model; data for all citers are shown. **c,d** The same data as those in panel **b** but now separated by the racial/ethnic category of the citer: **c** White citers and **d** citers of color. **e** The over-/under-citation of different author categories compared to their expected proportions under the relevant paper characteristics model; data for all citers are shown. **f,g** The same data as those in panel **e** but now separated by the racial/ethnic category of the citer: **f** White citers and **g** citers of color.

**Extended Data Figure 5.**
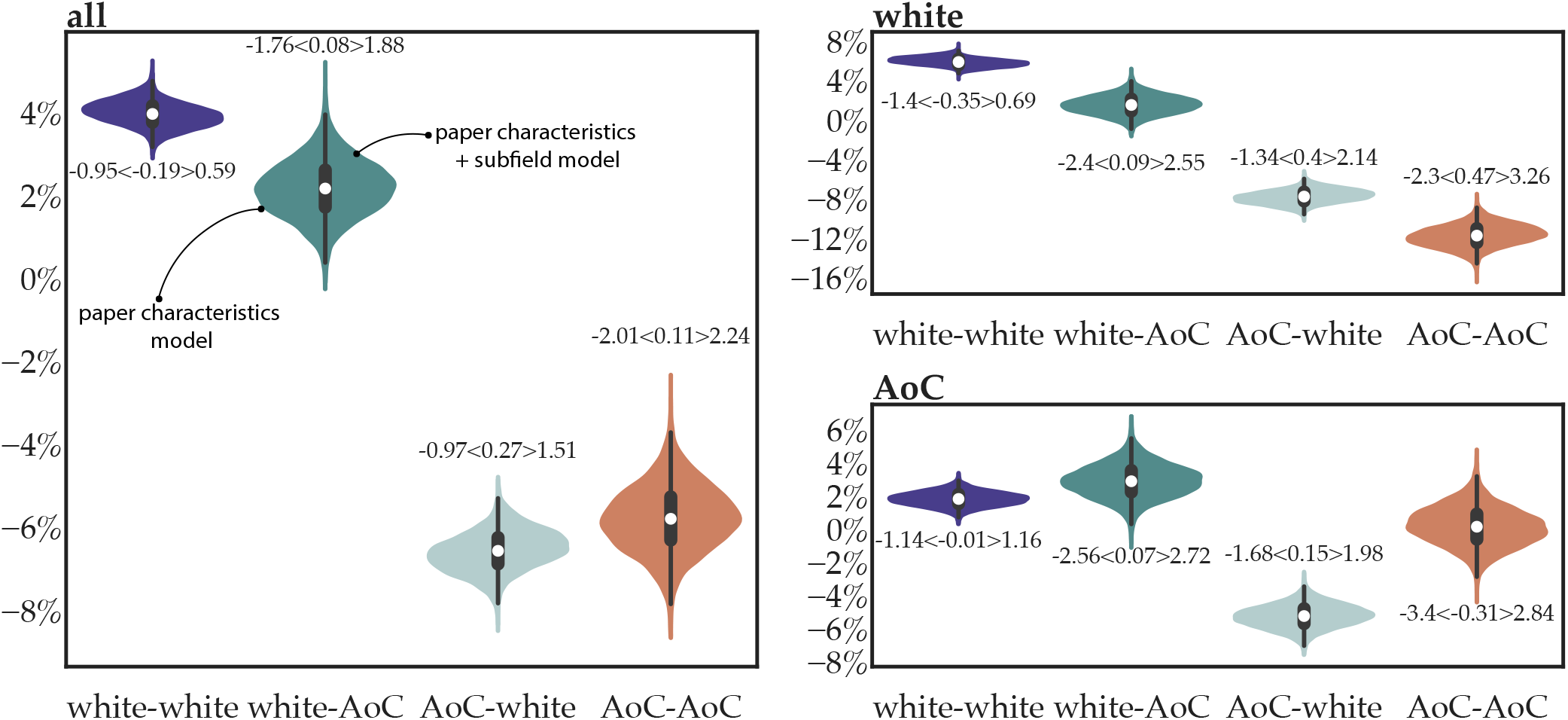
Effect of research subfields. Although the role of neuroscience subfields is not directly accounted for within the primary analyses, it is important to understand whether and to what extent relationships between gender and subfield confound our results. To assess this possibility, we conducted a sensitivity analysis on a subset of *Journal of Neuroscience* papers. The *Journal of Neuroscience* was chosen because it contains the most articles within our dataset, and because it uses a consistent sub-disciplinary classification scheme for its papers. Specifically, almost all of its papers are classified as either behavioral/systems, systems/circuits, neurobiology of disease, development/plasticity/repair, behavioral/systems/cognitive, or cellular/molecular. We fit two separate models predicting author race/ethnicity to the subset of 30555 articles with one of these classifications. The first model was the paper characteristics model with the same 6 factors described in the main text. The second model included those same 6 factors and also a 7-th factor: the subfield classification. Estimates of the over/under-citation of author race/ethnicity within these 30555 articles were then calculated from each of the two models. Here, the left side of the violin displays estimated over/under-citation under the paper characteristics model, and the right side of the violin displays estimated over/under-citation under the augmented paper characteristics model that also accounts for subfields. We show the 95% confidence intervals of the difference between the paper characteristics + sub-field model and the paper characteristics model. The results suggest that subfields likely have little impact on either the extent of citation imbalance or the discrepancy in citation behavior across citing author race/ethnicity.

**Extended Data Figure 6.**
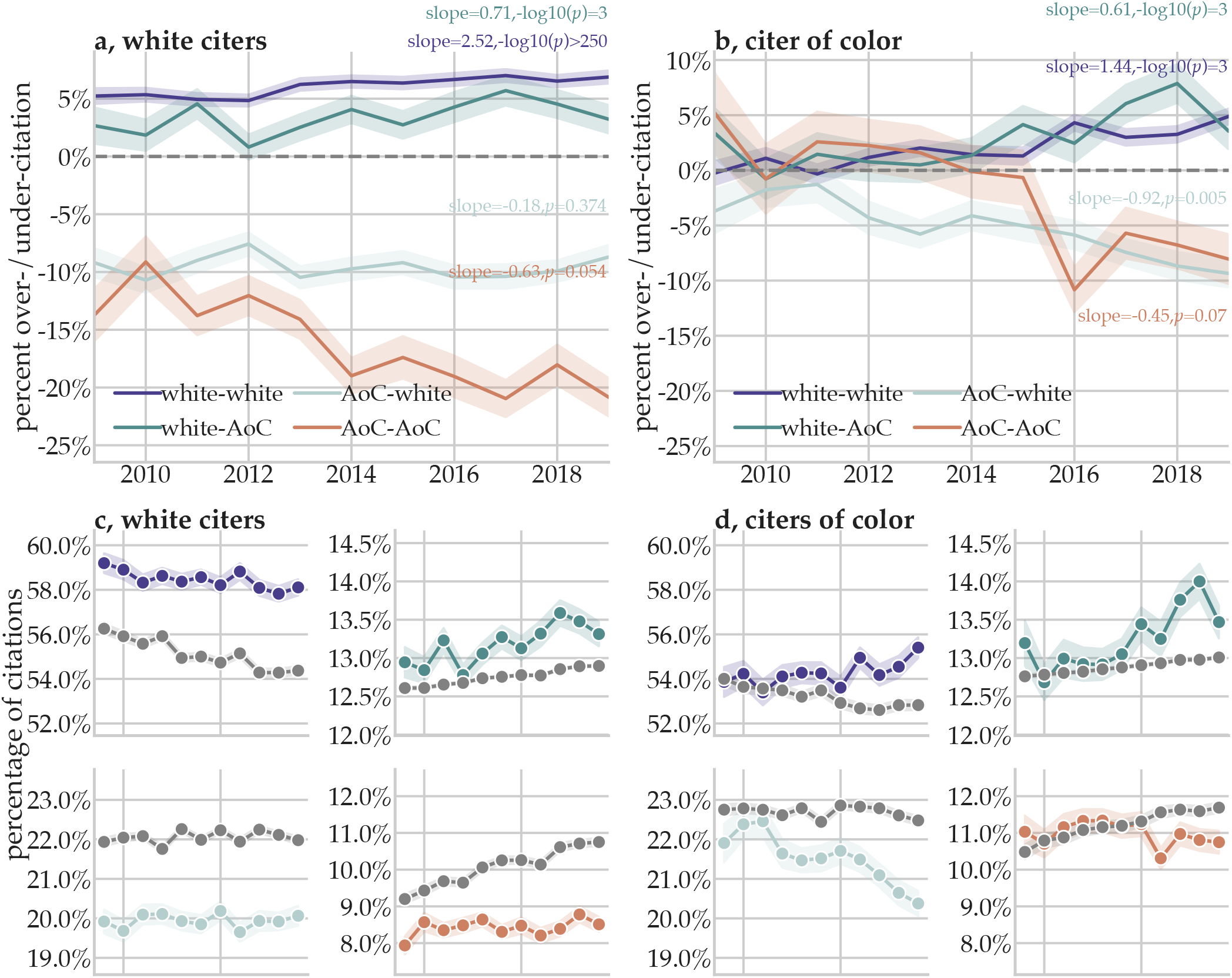
Temporal trends of over/under-citation (Census Model). **a,b** The extent of over/under-citation across author racial/ethnic categories as a function of time, within WW (**a**) and C∪C (**b**) reference lists. The line represents over/under-citation within the literature in a given year. Shaded regions represent the 95% confidence interval of each over/under-citation estimate, calculated from 1,000 bootstrap resampling iterations. **c,d** Observed (colored) and expected (grey) citation proportions within WW (**c**) reference lists and C∪C (**d**) reference lists. Within each section, we show the observed and expected proportion of citations given by that group to WW papers (*top left*), WC papers (*top right*), CW papers (*bottom left*), and CC papers (*bottom right*). Points represent the estimated citation percentage as a function of year; shaded regions represent the 95% confidence interval of the observed citation rate, calculated from 1,000 bootstrap resampling iterations.

**Extended Data Figure 7.**
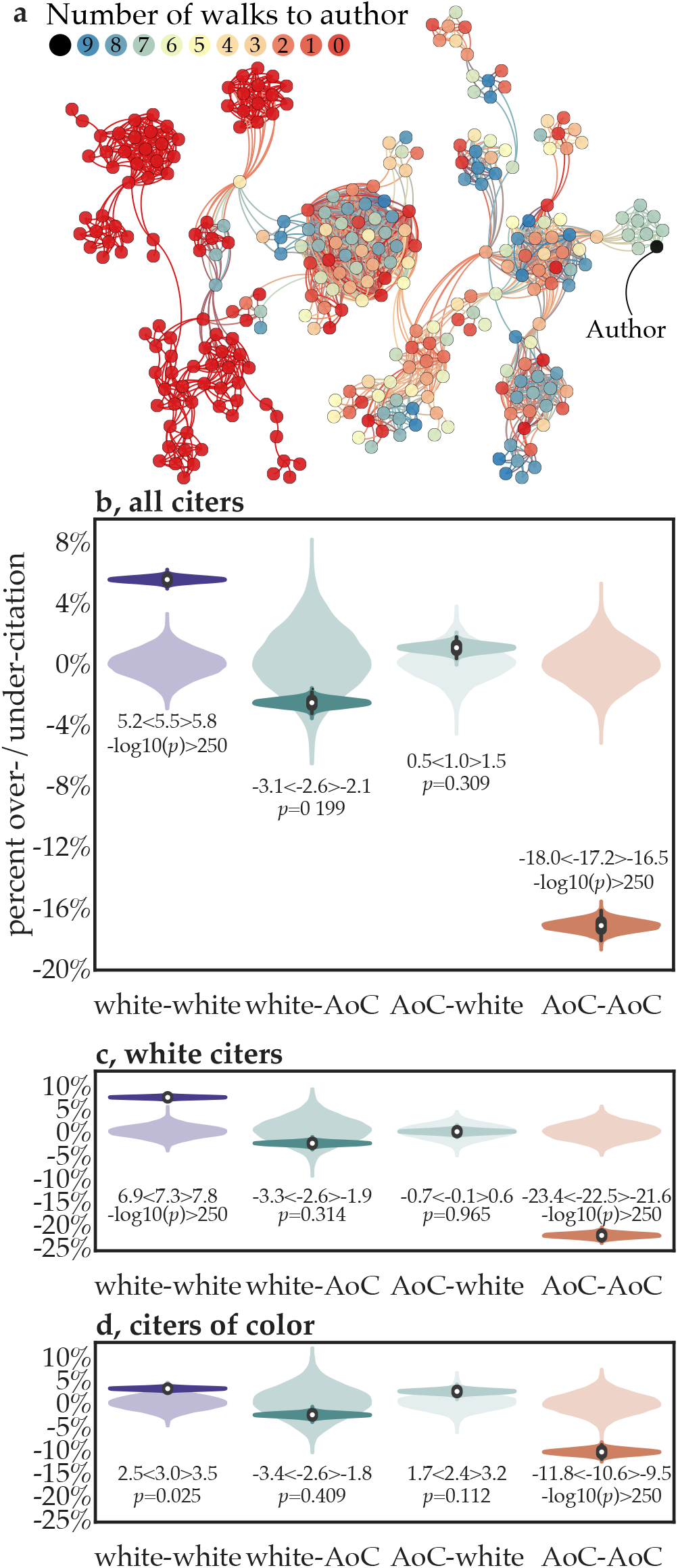
Estimating over/under-citation of papers based on random walks between authors in a co-authorship network. In the random walker model, we send random walkers out from the citing authors, allowing them to land on random citations. For each paper, we generated a custom co-authorship network that only contained connections between authors who had co-authored a paper prior to the citing paper we are considering. **a** The number of times a random walker visits each node, starting from the citing author node in black on the far left. Here, the lengths of each walk are set to the maximum distance between the citing author and all other authors. The chance of a random walker reaching an author is influenced by the network structure; for example, a random walker can become “stuck” in a group of highly connected authors. **b** The over/under-citation of different author racial/ethnic groups compared to their expected proportions under the random walk model. Results for **c** White citers and **d** citers of color. White authors are significantly over-cited and papers by two authors of color are significantly under-cited. This effect was driven more by White authors than by authors of color. When considering intersectionality, there is a 39 (95%CI=2.56,86.06) percentage point gap between men CC papers and women CC papers. There is a 33.23 (95%CI=1.98,60.82) percentage point gap between White men and men CC papers. There is a 69.08 (95%CI=67.39,70.55) percentage point gap between White men papers and White women papers. There is a 2.76 (95%CI=-19.5,18.44) percentage point gap between White women papers and women CC papers. Finally, there is a 35.85 (95%CI=-68.0,-7.98) percentage point gap between men CC papers and White women papers.

**Extended Data Figure 8.**
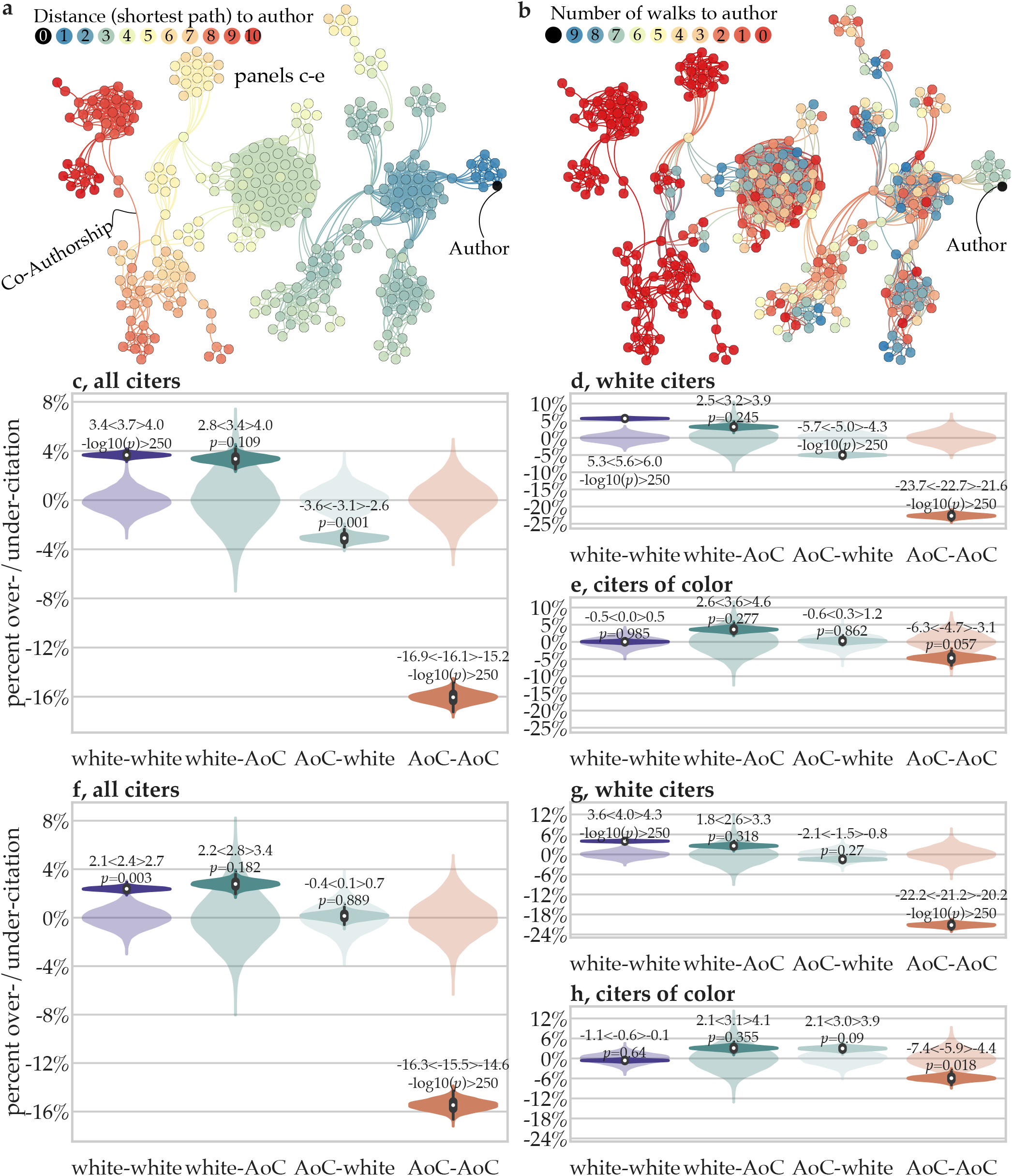
Estimating over/under-citation of papers based on author race/ethnicity when modeling citation practices on co-authorship networks (Census Model). **a** Shortest path distance and **b** the number of times a random walker visits each node, starting from the citing author node in black on the far left. These two models are similar, in that they both measure how close authors are to other authors in the co-authorship network. However, as can be seen in, whereas (i) the path length between two authors and (ii) the chance of a random walker reaching an author from another author are similar, the latter is more highly influenced by the structure of the network. For example, a random walker can become “stuck” in a group of highly connected authors. **c** The over/under-citation of different author racial/ethnic categories compared to their expected proportions under the shortest path distance model; data from all citers are shown. The same data as that shown in panel **c**, but now plotted separately for **d** White citers and **e** citers of color. The over/under-citation of different author racial/ethnic categories compared to their expected proportions under the random walks model; data from all citers are shown. The same data as that shown in panel **e**, but now plotted separately for **g** White citers and **h** citers of color.

**Extended Data Figure 9.**
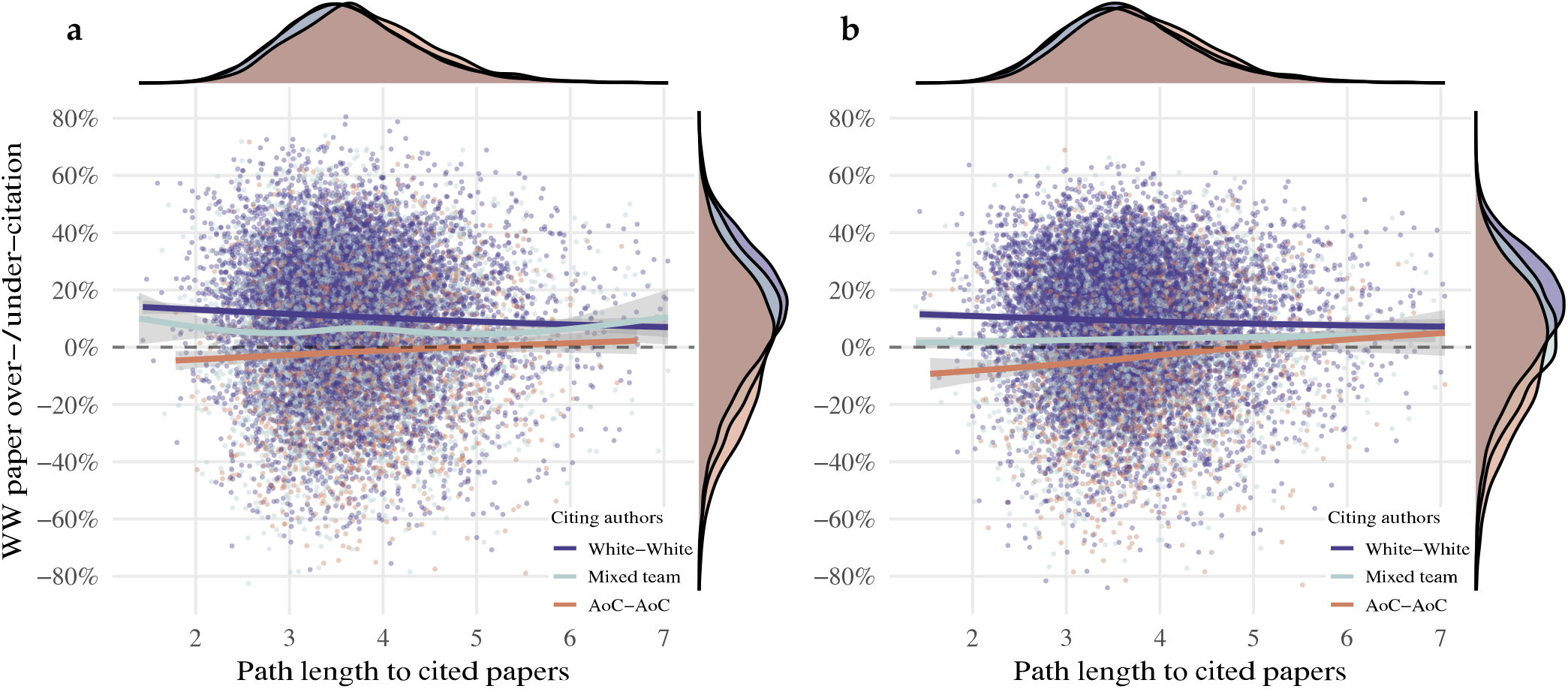
Relationships between authors’ path length to cited papers and racial/ethnic citation imbalance. Panel **a** shows results using the Florida model and panel **b** shows results using the Census model. In both panels, points represent individual papers. The value along the x-axis gives the average path length along the co-authorship network from citing paper authors to cited paper authors; the value along the y-axis gives the percent over- or under-citation of WW papers within a given reference list. Points are colored by the race/ethnicity of the papers’ first and last authors. Overlaid lines give the smoothed association between path length and citation behavior for each race/ethnicity group. Trend lines are estimated using a generalized additive model with cubic spline basis functions. Appended density plots show the marginal distributions of path length to cited papers (*top*) and WW paper over-/under-citation (*right*).

**Extended Data Figure 10.**
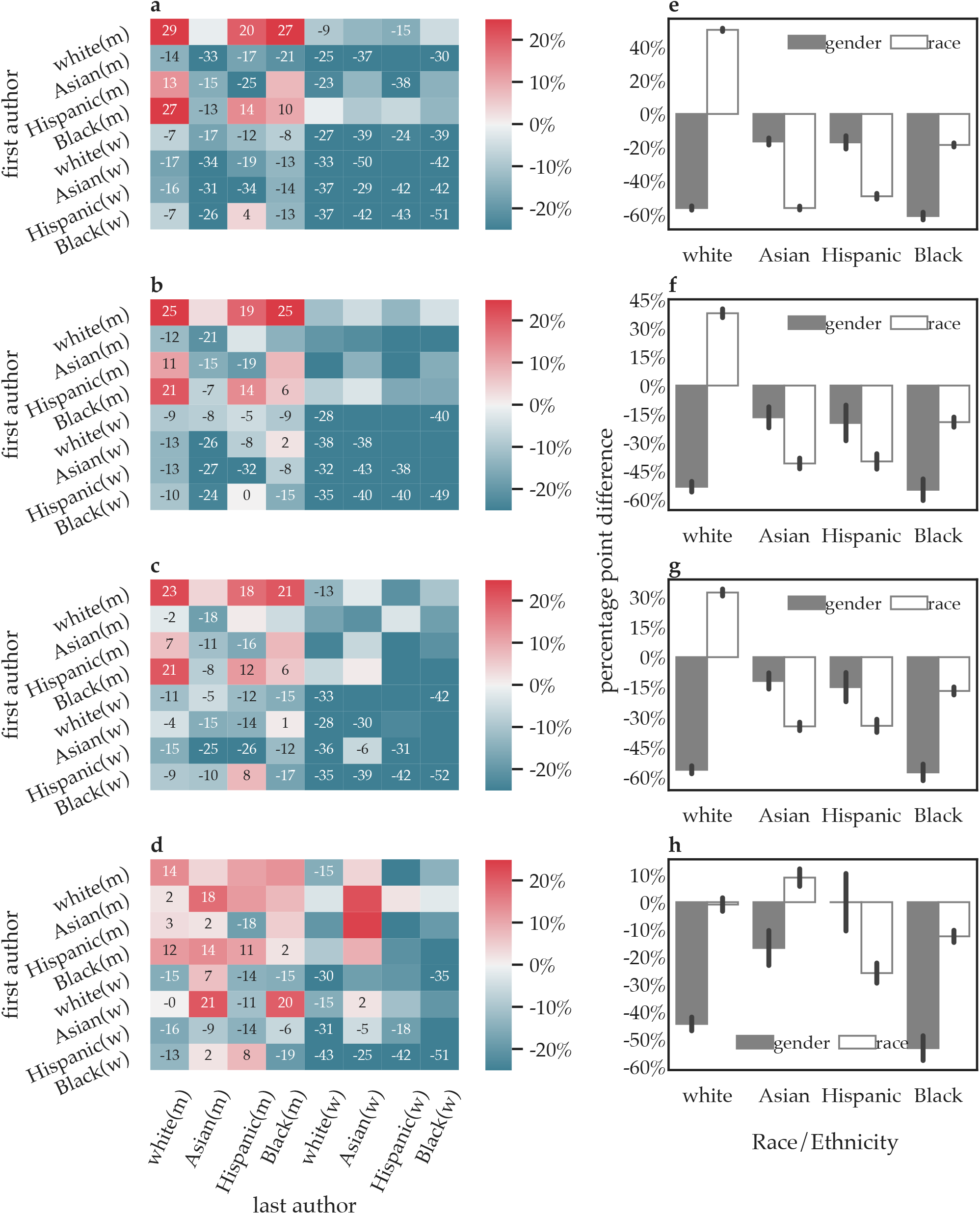
Citation costs at the intersection of gender and race/ethnicity (Random draws model; data separated by racial/ethnic category of *citers*). (**a-d**) We calculated the over/under-citation of papers for each combination of first author racial/ethnic and gender categories (y-axis) and last author racial/ethnic and gender categories (x-axis). Here, we compare the observed citation rates for each racial/ethnic and gender category to those that would be expected if base rates were defined by the random draws model for WW citers (**a**), CW citers (**b**), WC citers (**c**), and CC citers (**d**). We generate a null model of expected over/under-citation rates, where the expected citation counts are the probabilities from the random draws model that account for expected citation rates, and the observed citations counts are generated by pseudo-randomly citing based on those probabilities. For statistical inference, *p*-values are then calculated using this null distribution, and values that pass Holm-Bonferroni correction at *p* = 0.05 are annotated. (**e-h**) Next, we plot the percentage point difference in over/under-citation, comparing men and women authors within each race/ethnicity (*gender*; grey), and White authors to Asian, Black, and Hispanic authors (*race*; white), separately for WW citers (**e**), CW citers (**f**), WC citers (**g**), and CC citers (**h**).

**Extended Data Figure 11.**
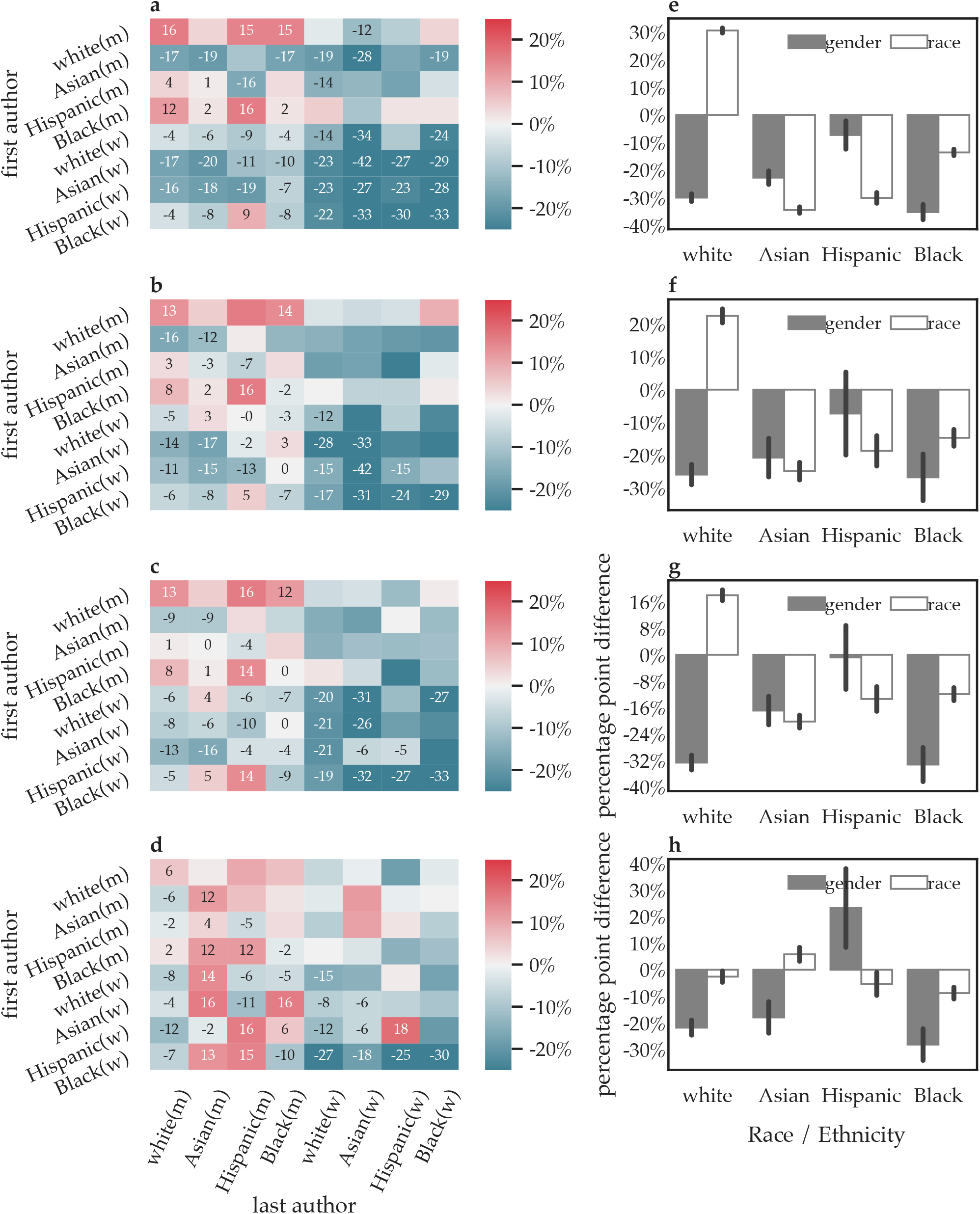
Citation costs at the intersection of gender and race/ethnicity (Relevant paper characteristics model; data separated by racial/ethnic category of *citers*). (**a-d**) We calculated the over/under-citation of papers for each combination of first author racial/ethnic and gender categories (y-axis) and last author racial/ethnic and gender categories (x-axis). Here, we compare the observed citation rates for each racial/ethnic and gender category to those that would be expected if base rates were defined by the paper characteristics model for WW citers (**a**), CW citers (**b**), WC citers (**c**), and CC citers (**d**). We generate a null model of expected over/under-citation rates, where the expected citation counts are the probabilities from the relevant paper characteristics model that account for expected citation rates, and the observed citations counts are generated by pseudo-randomly citing based on those probabilities. For statistical inference, *p*-values are then calculated using this null distribution, and values that pass Holm-Bonferroni correction at *p* = 0.05 are annotated. (**e-h**) Next, we plot the percentage point difference in over/under-citation, comparing men and women authors within each race/ethnicity (*gender*; grey), and White authors to Asian, Black, and Hispanic authors (*race*; white), separately for WW citers (**e**), CW citers (**f**), WC citers (**g**), and CC citers (**h**).

**Extended Data Figure 12.**
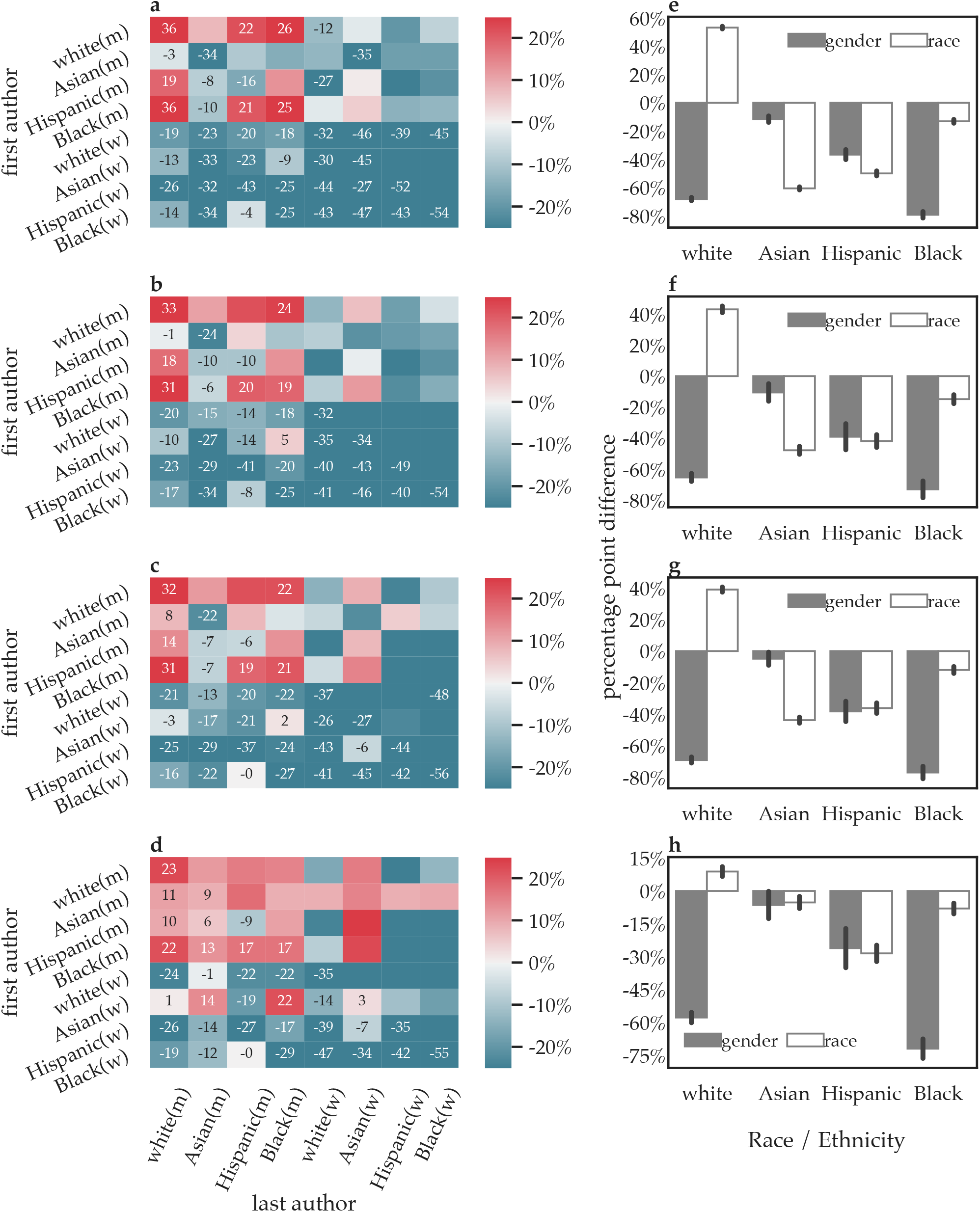
Citation costs at the intersection of gender and race/ethnicity (Shortest paths distance model; data separated by racial/ethnic category of *citers*). (**a-d**) We calculated the over/under-citation of papers for each combination of first author racial/ethnic and gender categories (y-axis) and last author racial/ethnic and gender categories (x-axis). Here, we compare the observed citation rates for each racial/ethnic and gender category to those that would be expected if base rates were defined by the co-authorship network distance model for WW citers (**a**), CW citers (**b**), WC citers (**c**), and CC citers (**d**). We generate a null model of expected over/undercitation rates, where the expected citation counts are the probabilities from the shortest paths distance model that account for expected citation rates, and the observed citations counts are generated by pseudo-randomly citing based on those probabilities. For statistical inference, *p*-values are then calculated using this null distribution, and values that pass Holm-Bonferroni correction at *p* = 0.05 are annotated. (**e-h**) Next, we plot the percentage point difference in over/under-citation, comparing men and women authors within each race/ethnicity (*gender*; grey), and White authors to Asian, Black, and Hispanic authors (*race*; white), separately for WW citers (**e**), CW citers (**f**), WC citers (**g**), and CC citers (**h**).

**Extended Data Figure 13.**
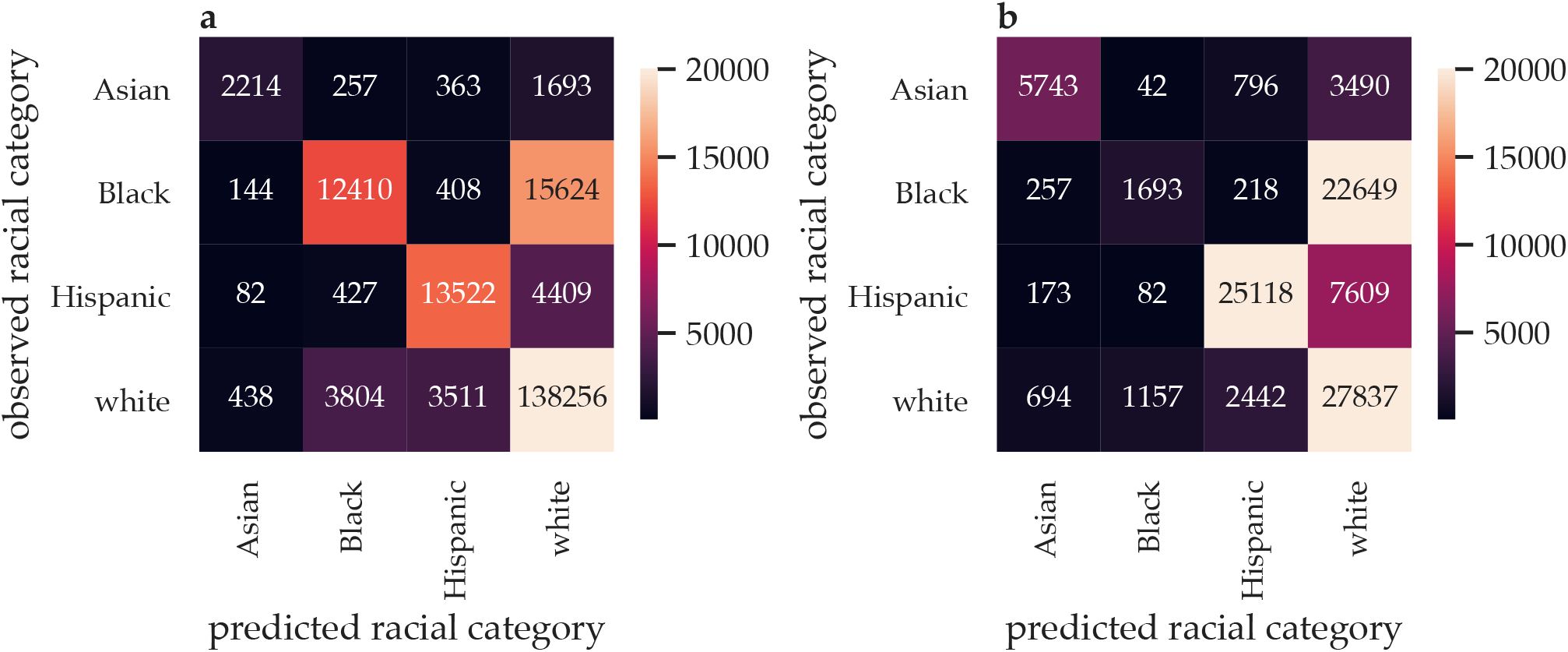
Confusion matrices for the Florida and Census models. **a** The ethnicolor Florida model confusion matrix demonstrating the accuracy of predictions; see the generally lighter colors along the diagonal and darker colors off the diagonal. **b** The ethnicolor Census model. In contrast to the Florida model, note that here many Black and Hispanic people are inaccurately predicted to be White. Data from https://github.com/appeler/ethnicolr.

